# TRIM62 promotes lipid raft-mediated entry of influenza A virus by regulating WASH localization

**DOI:** 10.1101/2025.09.28.679027

**Authors:** Kajal Gupta, Sampurna Pal, Roohani Bajaj, Tejal Pathak, Gaganpreet Kaur, Indranil Banerjee

## Abstract

Viruses employ diverse strategies to gain entry into cells to reach their replication sites. Influenza A virus (IAV), a respiratory pathogen, takes advantage of clathrin-dependent endocytosis or macropinocytosis for its internalization. Here, we report that TRIM62, a member of the tripartite motif (TRIM) protein family, is required for IAV entry, independent of its E3 ubiquitin ligase activity. We find that TRIM62 specifically functions in a clathrin-independent, lipid raft-mediated pathway, which IAV exploits to enter the endocytic network. Additionally, we reveal the involvement of the WASH complex and the retromer component VPS35 in this pathway beyond their canonical functions in endosomal sorting. We observe that a pool of WASH and VPS35 localize to the plasma membrane and associate with lipid rafts, in addition to their typical endosomal presence. While WASH subunits except FAM21 play a proviral role in IAV endocytosis and intracellular trafficking, VPS35 acts antagonistically. We show that TRIM62 counteracts the antiviral function of VPS35 by limiting FAM21-VPS35 interaction. By directly binding WASH in the cytosol, TRIM62 restricts its VPS35-mediated endosomal recruitment and thereby enhances its surface availability to facilitate IAV entry. Together, this study uncovers previously unrecognized roles of TRIM62 and endosomal sorting machinery in IAV entry, offering new antiviral targets.

## Introduction

Owing to its limited coding capacity, IAV has evolved elegant strategies to manipulate and co-opt host proteins and vesicular trafficking pathways for its entry. Given the centrality of vesicular trafficking to IAV entry, spanning from its initial endocytic uptake to fusion at the late endosome, a detailed understanding of how the virus takes advantage of this essential cellular process can reveal fundamental mechanistic insights into viral entry and illuminate potential host targets for therapeutic intervention.

The starting point for this study was the identification of TRIM62 as a host mediator of IAV entry, revealed through our focused RNAi screens targeting twenty tripartite motif (TRIM)-containing proteins that had been implicated in previous CRISPR-Cas screens against IAV infection^1–9^. The TRIM protein family in humans comprises over eighty distinct members, characterized by the presence of a conserved N-terminal RING domain with an E3 ubiquitin ligase activity, one or two B-box domains, a central coiled-coil domain (CC), and in some members, an additional C-terminal PRY-SPRY (P-S) domain^10^. Beyond their involvement in diverse cellular processes, several TRIM proteins are known to play critical roles in viral infections, with most functioning as antiviral factors^11^. However, our study revealed a contrasting proviral role of TRIM62, supporting IAV endocytosis and its downstream trafficking.

The unanticipated role of TRIM62 in IAV endocytosis prompted us to investigate its potential involvement in specific endocytic pathway(s). IAV primarily internalizes into host cells through clathrin-mediated endocytosis (CME)^12,13^, and less frequently via clathrin-independent endocytosis^14,15^, and macropinocytosis^16^. To examine which endocytic pathway(s) TRIM62 is involved in, we examined model cargoes specific to pathways including CME, macropinocytosis and lipid raft. We found that TRIM62 deficiency did not negatively impact the uptake of clathrin-dependent cargoes such as epidermal growth factor (EGF) and transferrin (Tfn), or macropinocytic cargo 70 kDa dextran (Dex). However, it markedly reduced the uptake of the lipid raft cargo cholera toxin B-subunit (CTxB), suggesting TRIM62’s involvement in the lipid raft-mediated pathway. CTxB binds GM1 glycosphingolipids and induces higher order clustering in the plasma membrane. It is often used as a reporter of lipid rafts enriched in sterols, sphingolipids and saturated phospholipids^17–19^. We observed that CTxB colocalizes with IAV both at the plasma membrane and during endocytosis, and reasoned that IAV can also internalize via a lipid raft pathway involving TRIM62, in addition to classical CME or macropinocytosis.

Since clathrin-independent, lipid raft pathways can involve caveolin, dynamin, Arp2/3-driven branched actin, and plasma membrane-associated nucleation-promoting factors (NPFs) WAVE and N-WASP ^19–21^, we examined whether IAV co-opts these factors for its entry and found that none facilitated viral uptake. We also examined the endosome-associated NPF WASH (Wiskott-Aldrich Syndrome protein and SCAR Homolog), as it was found to be recruited to the plasma membrane during *Salmonella* invasion^22^ and to facilitate clathrin-independent endocytosis of human papillomavirus (HPV)^23^. We found that silencing WASH1, the WASH complex subunit with NPF function, significantly reduced IAV internalization, similar to the effect of TRIM62 depletion, underscoring WASH1’s critical role in viral entry. Strikingly, we also find that while WASH1’s presence is essential, its NPF activity is dispensable for viral entry, highlighting that its scaffolding rather than actin assembly function, drives this process.

The WASH complex is a pentamer of WASH1, CCDC53, SWIP, Strumpellin, and FAM21, of which the WASH1 subunit belongs to the WASP family and is responsible for Arp2/3 activation via its C-terminal verprolin homology sequence, connecting sequence, and acidic sequence (VCA) domain ^24–26^. The WASH complex is recruited to endosomes either via interaction of FAM21 with the retromer subunit VPS35 or through direct endosomal membrane binding of SWIP in a retromer-independent manner ^27^. We observed that WASH1 and VPS35 colocalize with the lipid raft marker CTxB at the plasma membrane and at endosomes, but strikingly, they play opposing roles in the internalization of CTxB and IAV: WASH1 depletion reduces the uptake of CTxB and IAV, whereas VPS35 deficiency enhances it, indicating a suppressive role for VPS35. Although the WASH regulatory complex (SHRC) comprising CCDC53, SWIP, Strumpellin, and FAM21, showed no role in CTxB uptake, the SHRC subunits differentially influenced IAV entry: FAM21 acted as an antiviral factor, whereas the others promoted viral entry, highlighting FAM21’s distinct role in this process. This highlights a shared lipid raft-dependent pathway, yet distinct endocytic requirements for CTxB and IAV. Given the involvement of TRIM62 and WASH1 in the lipid raft pathway and IAV internalization, we next explored whether they associate to cooperatively regulate lipid raft-mediated viral endocytosis. We found that TRIM62 colocalizes with the WASH complex at the cell surface and interacts with all its subunits, most notably through a direct interaction with FAM21, the subunit that restricts viral entry. However, TRIM62 showed no interaction with the endosomal retromer components, including VPS35, indicating its specific association with the WASH complex. Interestingly, TRIM62 deficiency led to an enhanced the FAM21-VPS35 interaction, leading to an increased recruitment of the WASH complex to endosomes. This highlights TRIM62’s regulatory role in WASH recruitment to endosomes and in modulating the antiviral FAM21-VPS35 axis. Notably, while TRIM62 or VPS35 depletion did not affect total WASH levels, it differentially regulated WASH spatial distribution, producing opposite outcomes for IAV entry: TRIM62 depletion caused WASH overabundance on endosomes and impaired viral entry, whereas VPS35 loss redistributed WASH to the cytosol and enhanced entry. Interestingly, co-depletion of TRIM62 with VPS35 fully restored viral entry to basal levels. However, viral entry was markedly reduced when WASH1 and VPS35 were simultaneously depleted. These findings suggest that TRIM62 and VPS35 function antagonistically in regulating WASH’s endosome-cytosol dynamics, thereby modulating viral entry, with WASH1 functioning as a critical entry mediator.

Together, our study uncovers a noncanonical, NPF activity-independent role of the WASH complex in lipid raft-dependent entry of IAV. In this pathway, TRIM62 and VPS35 engage in reciprocal regulation on WASH localization, thereby maintaining its homoeostasis and enabling productive viral entry.

## Results

### Focused RNAi screening of TRIM family proteins identifies TRIM62 as an IAV entry mediator

To revalidate the involvement of TRIM proteins identified as hits in previous CRISPR-Cas screens against IAV infection^1–9^, we conducted a focused RNAi screen, targeting genes that were identified in at least two independent screens. We included scrambled siRNA (siControl) as a negative control and siRNA targeting the ATP6V1B2 subunit of vacuolar H^+^-ATPase or v-ATPase (siATP6V1B2) as a positive control. We also included bafilomycin A1 (BafA1), a potent v-ATPase inhibitor, as an additional positive control. Using viral nucleoprotein (NP) expression as a readout for infection, the screen identified fifteen TRIM proteins whose depletion significantly reduced infection (**Fig. 1a**). Since viral gene expression initiates only after the viral ribonucleoprotein complexes (vRNPs) of incoming viruses are imported into the nucleus, the final step of viral entry (**Fig. 1b**), we addressed whether any of the identified proviral TRIM proteins are involved in viral entry. We conducted a subsequent RNAi screen targeting these fifteen TRIM proteins for vRNP nuclear import. Here, we included an siRNA targeting karyopherin (importin) β-1 (siKPNB1) and importazole (Ipz), an importin β inhibitor, as positive controls, along with siATP6V1B2. In this screen for vRNP nuclear import, we identified TRIM9, 35 and 62 as hits (**Fig. 1c**). Among them, TRIM62 depletion most effectively attenuated vRNP nuclear import, making it the strongest candidate for further investigation.

**Fig. 1.**
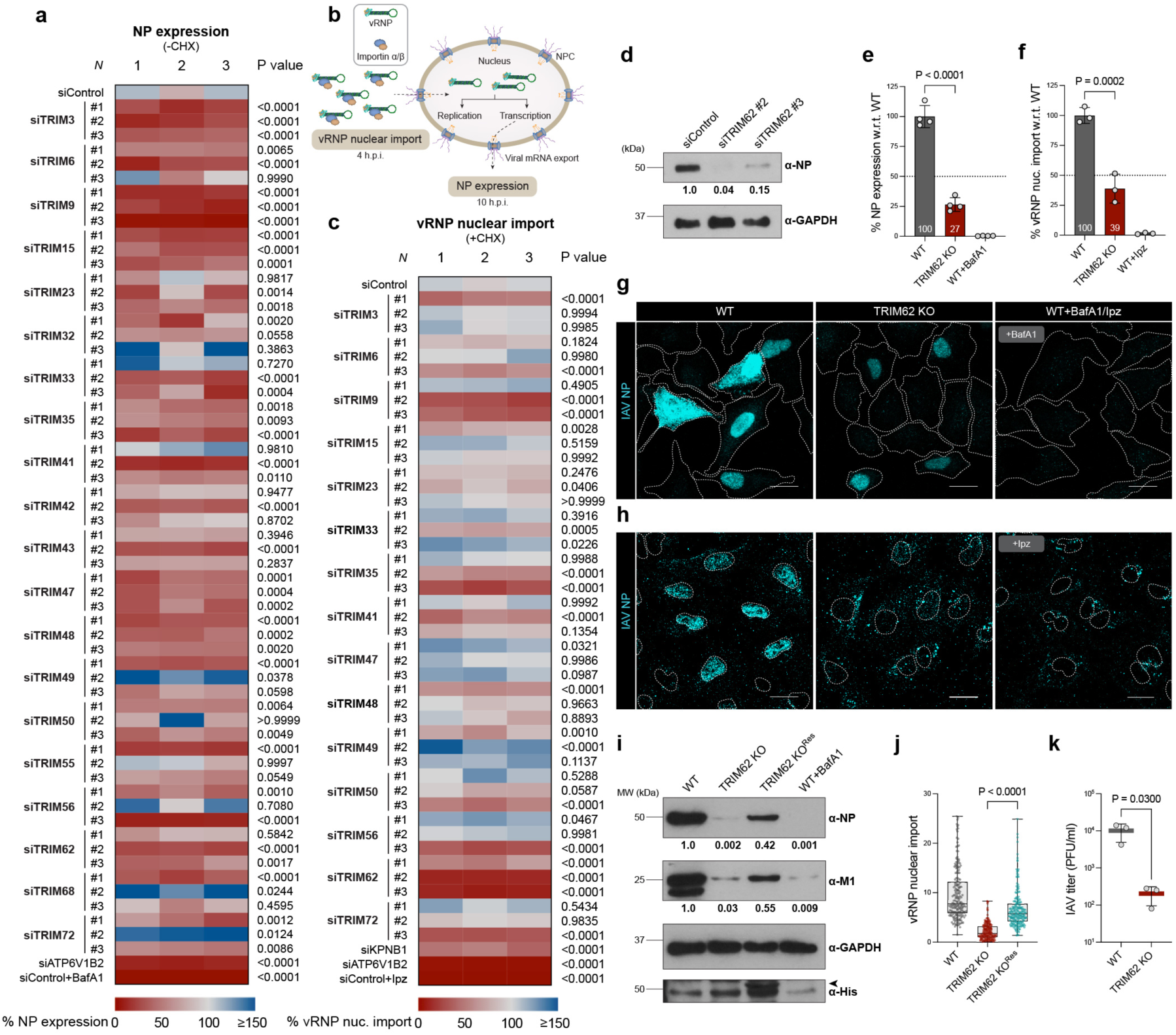
TRIM62 is required for IAV host cell entry. **a** Heatmap showing IAV infection in A549 cells infected with the X-31 (H3N2) strain (MOI = 0.01) in an siRNA screen targeting twenty TRIM proteins. A549 cells were treated with siControl or siATP6V1B2 targeting the v-ATPase complex. Additionally, the v-ATPase inhibitor bafilomycin A1 (BafA1) was used to block endosomal acidification, and thus infection. The readout for infection was % NP expression at 10 h.p.i. (NP was detected by indirect immunofluorescence, IIF). The screen was repeated three times, and infection inhibition by three independent siRNAs are shown along with corresponding P values. *n* > 2000 cells per sample. **b** Schematic of vRNP nuclear import and subsequent NP expression during IAV infection. **c** Heatmap showing normalized values of % vRNP nuclear import at 4 h.p.i. in an siRNA screen targeting fifteen TRIM proteins identified as supportive factors for IAV infection. A549 cells were treated with siControl or siATP6V1B2 or siKPNB1 targeting the importin β1 gene or importazole (Ipz), an inhibitor that blocks importin β. Cycloheximide (CHX) was added to prevent the synthesis of new viral proteins. NP was detected by IIF and the percentage of NP-positive nuclei was calculated as a readout for vRNP nuclear import. The screen was repeated three times, and vRNP nuclear import inhibition by three independent siRNAs are shown along with corresponding P values. *n* > 800 cells per sample. **d** Western blot showing reduced NP levels at 10 h.p.i. in TRIM62-depleted cells compared to siControl. GAPDH was used as a loading control. **e** Quantification showing reduced NP expression at 10 h.p.i. in TRIM62 knockout (KO) cells compared to WT cells. *n* > 5500 cells per sample. **f** Quantification showing reduced vRNP nuclear import at 4 h.p.i. in TRIM62 KO cells compared to WT cells. *n* > 1100 cells per sample. **g,h** Representative confocal microscopy images showing NP (cyan) for IAV infection (**g**) and vRNP nuclear import (**h**). Cell and nuclear boundaries are outlined. Scale bars, 20 µm. **i** Western blots showing restored viral NP and M1 levels at 10 h.p.i. in TRIM62 rescue cells compared to TRIM62 KO cells. Arrow marks the specific TRIM62-His band. GAPDH was used as a loading control. **j** Quantification showing restored vRNP import (NP intensity in nuclei) at 4 h.p.i. in TRIM62 rescue cells compared to TRIM62 KO cells. *n* = 200 cells per sample. **k** Quantification of IAV titer by viral plaque assay. PFU, plaque-forming unit. Data in **a**,**c**,**e**,**f** were analyzed using one-way ANOVA using multiple-comparisons. Data in **j**,**k** were analyzed using unpaired two-tailed *t*-test, P values are indicated. P values < 0.05 were considered significant. All bar graphs show the mean of n = 3 ± SD, and all data are representative of three biological replicates (N = 3).

Next, we revalidated TRIM62’s role in viral gene expression by western blotting and indirect immunofluorescence (IIF), which showed a marked reduction in NP expression upon silencing the gene with two independent siRNAs (**Fig. 1d, Supplementary Fig. 1a**). We observed similar results for vRNP nuclear import, as evidenced by IIF (**Supplementary Fig. 1b**). Due to the unavailability of a specific antibody that can detect endogenous TRIM62, we assessed the knockdown (KD) efficiency of the siRNAs by reverse transcription polymerase chain reaction (RT-PCR) (**Supplementary Fig. 1c**). Since siTRIM62 #2 demonstrated higher KD efficiency than the other, we used it (hereafter referred to as siTRIM62) for all subsequent RNAi-based experiments. Next, we addressed whether TRIM62’s role in IAV infection is specific to viral strain or cell line by examining multiple strains including WSN (H1N1, spherical), NYMC (H3N2, spherical) and Udorn (H3N2, filamentous), along with HeLa, HepG2 and Huh-7 cells. TRIM62 deficiency significantly reduced infection in all cases, highlighting its general role (**Supplementary Fig. 1d-k**). However, subsequent experiments were carried out in A549 cells, as this lung-origin cell line closely recapitulates the virus’s natural target cells. Following RNAi results, we generated TRIM62 knockout (KO) in A549 cells using CRISPR-Cas9, and validated the KO cells by sequencing the gRNA-targeted genomic region and RT-PCR (**Supplementary Fig. 2a, b**). Although sequencing results confirmed successful editing at the target site, the genome-edited cells still retained about 13% of TRIM62 transcripts relative to wild-type (WT) cells. Consistent with TRIM62 KD phenotypes, the KO cells also reduced IAV (X-31) infection and vRNP nuclear import by 73% and 61%, respectively, compared to WT cells (**Fig. 1e-h**). Also, TRIM62 KO cells showed significantly reduced infection across other IAV strains including WSN, NYMC, and Udorn (**Supplementary Fig. 2c-f**). To further confirm TRIM62’s involvement, we performed rescue experiments by stably expressing His-tagged TRIM62 (TRIM62-His) in KO background. TRIM62-His-complemented KO cells showed restored IAV infection and vRNP nuclear import compared to KO cells (**Fig. 1i, j**). Viral plaque assay with supernatants from infected A549 cells (supernatants collected at 24 hours post-infection, h.p.i.) showed that TRIM62 KO cells produced nearly 50-fold fewer infectious IAV particles than parental cells (**Fig. 1k**). Together, we establish TRIM62 as a pro-viral host factor that supports IAV infection by facilitating vRNP nuclear import during entry.

### TRIM62 facilitates IAV endocytosis and downstream trafficking

After identifying TRIM62’s role in vRNP nuclear import, we investigated its involvement at earlier stages of viral entry. To this end, we performed quantitative high-content, imaging-based IAV entry assays that we had previously developed^28^, to examine TRIM62’s involvement at each sequential entry step including virion binding, endocytosis, HA acidification, fusion, and M1 uncoating. The sequential entry steps of IAV involving vesicular trafficking i.e. prior to vRNP nuclear import is shown in the schematic (**Fig. 2a**). Although virion binding to the plasma membrane was unaffected in TRIM62-deficient cells (**Fig. 2b, g, Supplementary Fig. 3a, b**), viral endocytosis (30 min post-infection) was reduced by 43% and 55% in TRIM62 KO and KD cells, respectively (**Fig. 2c, h, Supplementary Fig. 3c, d**). Consistent with the endocytosis defect, TRIM62-deficient cells showed an impairment at post-endocytic entry steps including HA acidification (1 h.p.i.), fusion (1.5 h.p.i.), and M1 uncoating (2 h.p.i.) (**Fig. 2d-f, i-k Supplementary Fig. 3e-j**). Thus, both KO and KD cells displayed comparable phenotypes across IAV entry steps. Strikingly, M1 uncoating (cytosolic M1 dispersal) exhibited the most pronounced phenotype among the entry steps, with 67% and 78% reductions in KO and KD cells (**Fig. 2f, k, and l**), respectively, compared to controls. Was this marked reduction in M1 uncoating (examined at 2 h.p.i.) due to delayed kinetics? Time-course analysis showed consistent impairment in uncoating, excluding this possibility (**Fig. 2m**). To further validate TRIM62’s involvement in uncoating, we stably overexpressed Myc-tagged TRIM62 in A549 cells and quantified cytosolic M1 intensity per unit area in TRIM62-overexpressing cells. Unlike percentage-averaged quantification, this method captured cell-to-cell differences and provided a more detailed measure of uncoating. Notably, TRIM62 overexpression significantly enhanced M1 uncoating, underscoring its key role in the process (**Fig. 2n**).

**Fig. 2.**
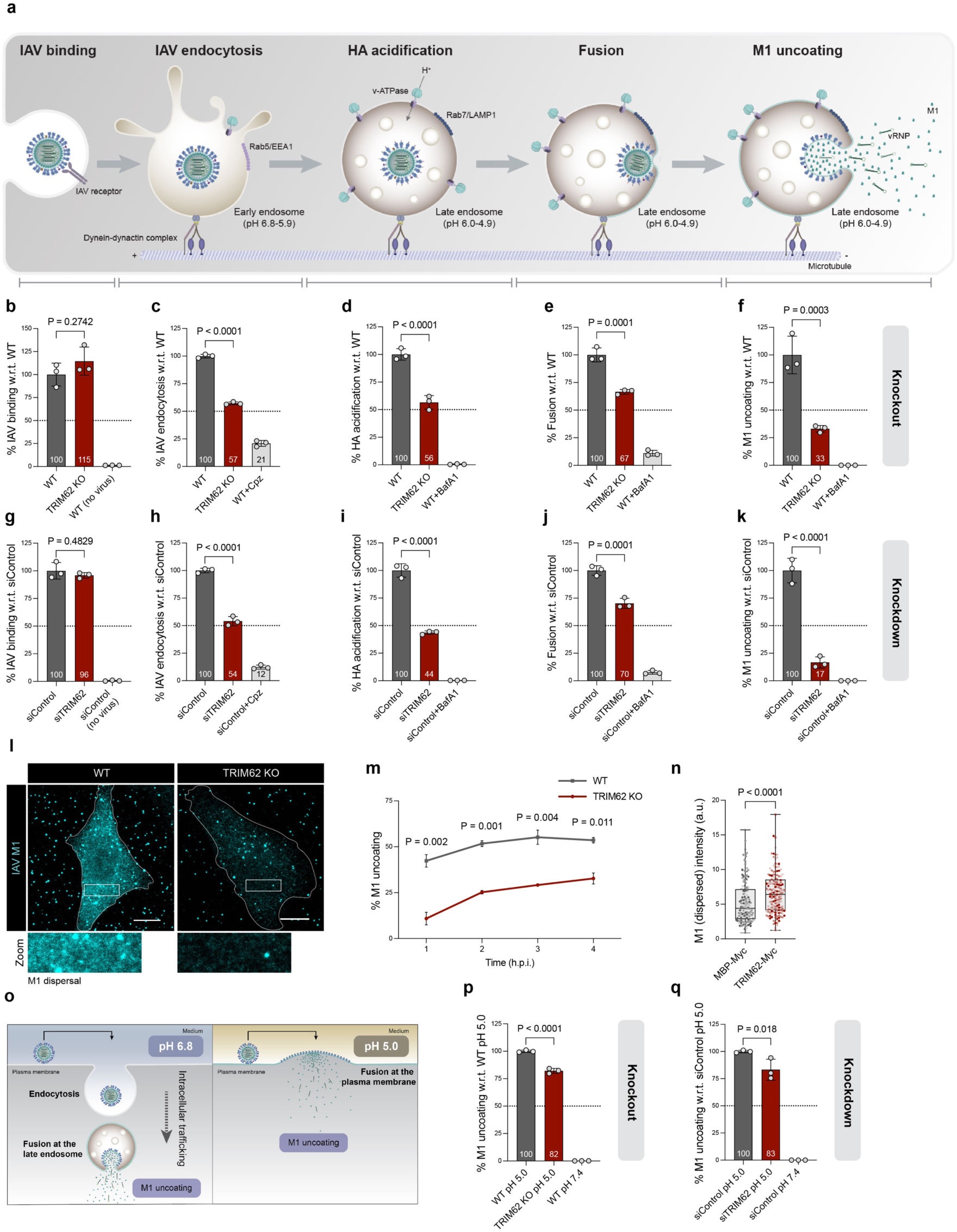
TRIM62 promotes IAV endocytosis and vesicular trafficking. **a** Schematic showing key events during IAV vesicular trafficking including IAV binding, IAV endocytosis, HA acidification, fusion, and M1 uncoating. **b-f** Quantification of percentages of IAV binding (**b**), endocytosis (**c**), HA acidification (**d**), Fusion (**e**), and M1 uncoating (**f**) at 0-, 30-, 60-, 90-, and 120-min post-infection, respectively, in WT and TRIM62 KO cells. Chlorpromazine (Cpz) was included as a positive control in IAV endocytosis assay, whereas BafA1 was used as a positive control for HA acidification, fusion, and M1 uncoating. *n* > 4000 cells/assay. **g-k** Quantification of percentages of IAV binding (**g**), IAV endocytosis (**h**), HA acidification (**i**), Fusion (**j**), and M1 uncoating (**k**) at 0, 30-, 60-, 90-, and 120-min post-infection, respectively in control and TRIM62 knockdown (KD) cells. *n* > 1500 cells per sample. **l** Representative confocal microscopy images of M1 uncoating, showing M1 (cyan). Insets show cytosolic M1. Scale bars, 10 µm. **m** Time-course analysis of M1 uncoating. Quantification of % M1 uncoating at 1, 2, 3, and 4 h.p.i. **n** Quantification of cytosolic M1 (dispersed) intensity in cells expressing MBP-Myc or TRIM62-Myc. *n* = 152 cells per sample. **o** Schematic showing plasma membrane-bypass M1 uncoating. IAV enters cells via endocytic routes at pH 6.8. When viral particles are induced to fuse at the plasma membrane under acidic condition (pH 5.0), the capsids disassemble, leading to M1 dispersal into the cytosol. **p,q** Quantification of plasma membrane-bypass M1 uncoating (% cells with dispersed M1) at pH 5 and pH 7.4 (negative control) in control and TRIM62 KO or KD cells. *n* > 1100 cells per sample. Data in **b-k,p,q** were analyzed using one-way ANOVA using multiple-comparisons. Data in **m,n** were analyzed using unpaired two-tailed *t*-test, P values are indicated. P values < 0.05 were considered significant. All bar graphs show the mean of n = 3 ± SD, and all data are representative of three biological replicates (N = 3).

Does TRIM62 directly facilitate M1 uncoating post-fusion, or is the uncoating defect in TRIM62-deficient cells a consequence of impaired vesicular trafficking? To clarify this, viral fusion was induced at the plasma membrane using a pH 5.0 medium, enabling direct capsid delivery to the cytosol, thereby bypassing endosomal entry^29^ (**Fig. 2o**). This led to only 18% and 17% reductions in uncoating in TRIM62 KO and KD cells, respectively (**Fig. 2p, q**). Although statistically significant, the moderate reductions in plasma membrane-bypass uncoating compared to the pronounced defects in uncoating through the endosomal route, highlight TRIM62’s primary involvement in vesicular trafficking. Collectively, these findings demonstrate that TRIM62 promotes IAV endocytosis and facilitates its vesicular trafficking to enable efficient uncoating.

### TRIM62 and WASH1 drive clathrin-independent, lipid raft-mediated IAV endocytosis

IAV exploits multiple endocytic pathways for entry, including clathrin-mediated endocytosis (CME) and non-clathrin pathways that include macropinocytosis and lipid raft-dependent pathways^12,14,16,30^, some of which are dynamin-dependent while others proceed independently of dynamin (**Fig. 3a**). Since we observed that TRIM62 deficiency impairs IAV endocytosis, we next investigated whether it selectively modulates specific endocytic pathway(s). To address this, we analyzed the cellular uptake of model endocytic pathway-specific cargoes in TRIM62 KD cells. For CME, we assessed EGF and Tfn uptake for 30 min and 10 min, respectively, with chlorpromazine (Cpz) and an siRNA targeting the clathrin heavy chain (siCLTC) serving as positive controls. Interestingly, while EGF internalization remained unaffected, Tfn uptake was significantly increased in TRIM62-deficient A549 cells (**Fig. 3b, c**). Although EGF is primarily a CME cargo, it can also enter cells through non-clathrin pathways^31^, whereas Tfn is a *bona fide* CME cargo. To further examine the effect of TRIM62 deficiency on CME, we assessed infection with dengue virus serotype 2 (DENV2), which enters HepG2 cells via CME^32^. Mirroring the Tfn uptake phenotype, DENV-2 infection also increased upon TRIM62 KD (**Supplementary Fig. 4a**), indicating that the IAV endocytosis defect in TRIM62-deficient cells was not due to impaired viral entry through CME.

**Fig. 3.**
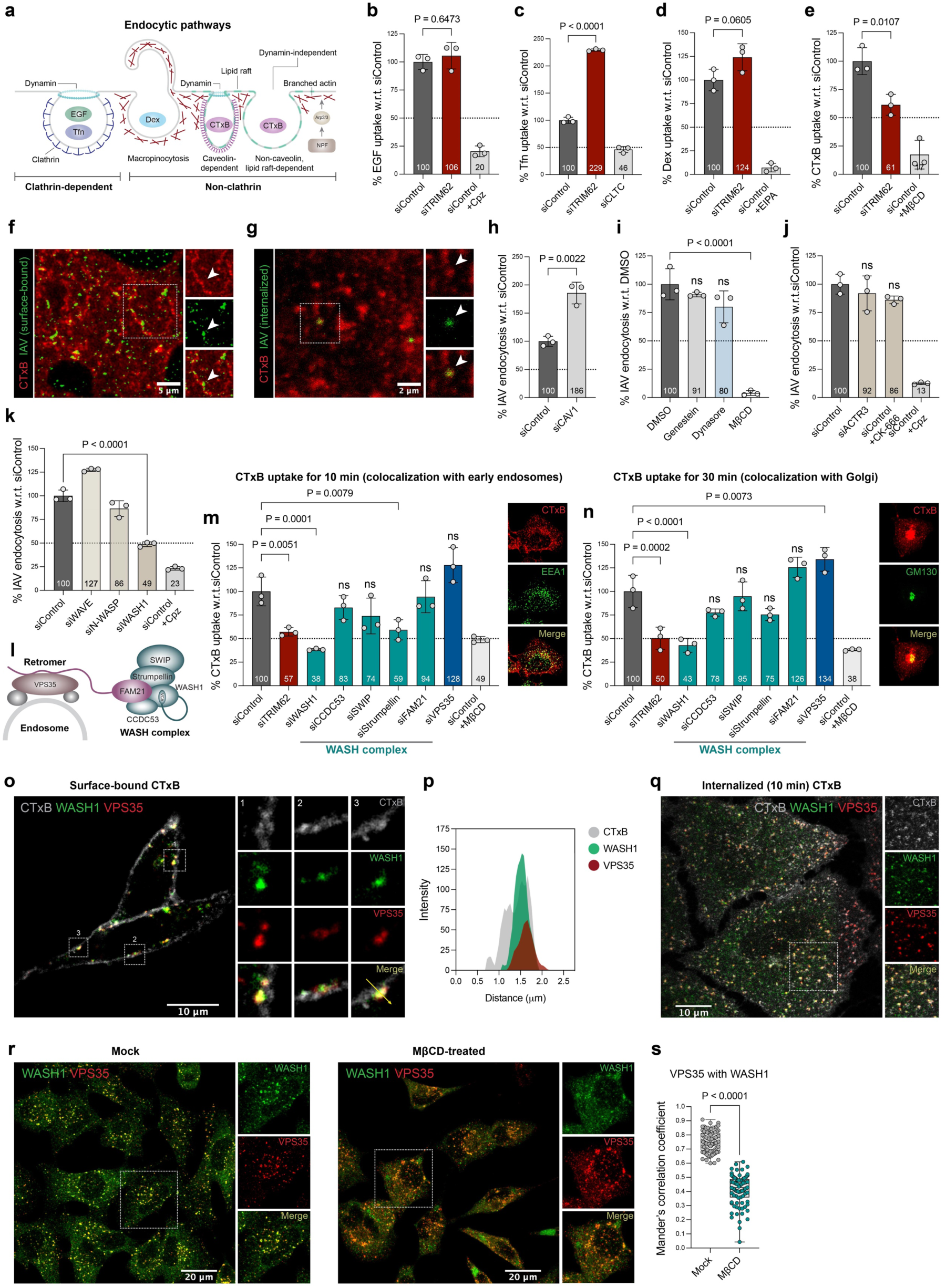
TRIM62 and WASH1 drive clathrin-independent, lipid raft–mediated IAV endocytosis. **a** Schematic showing cargo-specific clathrin-dependent and -independent pathways of endocytosis. Non-clathrin pathways include macropinocytosis, caveolin-dependent and -independent lipid raft-mediated endocytosis. **b-e** Quantification of EGF (**b**), Tfn (**c**), Dex (**d**), and CTxB (**e**) uptake in TRIM62-depleted cells, as compared to control cells. *n* ≥ 1000 cells per sample. **f** Confocal microscopy images showing surface-bound CTxB (red) and IAV (green). Insets show the distribution of CTxB and IAV on lipid rafts and white arrowheads indicate colocalization. Scale bars, 3 µm. **g** Confocal microscopy images showing CTxB and IAV 10 min post-internalization. Insets show the distribution of internalized CTxB and IAV and white arrowheads indicate colocalization. Scale bars, 2 µm. **h** Quantification of percentage of IAV endocytosis in control and CAV1 KD cells. *n* > 500 cells per sample. **i** Quantification of percentage of IAV endocytosis in DMSO-treated and inhibitor-treated cells. *n* > 500 cells per sample. **j** Quantification of percentage of IAV endocytosis in control cells, ACTR3 KD cells, and CK-666-treated cells. *n* ≥ 500 cells per sample. **k** Quantification of percentage of IAV endocytosis in control and WAVE, N-WASP, and WASH1 KD cells. *n* ≥ 500 cells per sample. **l** Schematic showing the recruitment of pentameric WASH complex via retromer at endosomal surface, specifically mediated by the interaction of FAM21 with VPS35. **m,n** Quantification of CTxB uptake in control and KD cells post 10 min (**m**) and 30 min (**n**) internalization. Images show CTxB uptake through colocalization of CTxB with EEA1+ endosomes at 10 min (**m**) and with GM130+ Golgi compartments at 30 min (**n**), respectively. *n* ≥ 1000 cells per sample. **o,p** Confocal microscopy images (**o**) showing surface-bound CTxB (grey), WASH1 (green) and VPS35 (red). Insets show the colocalization of CTxB, WASH1 and VPS35 at the plasma membrane. Scale bars, 10 µm. Fluorescence intensity profiles of CTxB, WASH1 and VPS35 along the yellow line are shown (**p**). **q** Confocal microscopy images showing CTxB, WASH1 and VPS35 10 min post internalization. Insets show the distribution of internalized CTxB, WASH1 and VPS35 and their colocalization. Scale bars, 10 µm. **r** Confocal microscopy images showing WASH1 (green) and VPS35 (red) in control and MβCD-treated cells. Insets show the distribution of WASH1 and VPS35. Scale bars, 20 µm. **s** Quantification of WASH1-VPS35 colocalization quantified by Mander’s correlation coefficient. *n* = 86 cells per sample. Data in **b-e,I,j,k,m,n** were analyzed using one-way ANOVA using multiple-comparisons. Data in **h,s** were analyzed using unpaired two-tailed *t*-test, P values are indicated. P values < 0.05 were considered significant. All bar graphs show the mean of n = 3 ± SD.

To determine TRIM62’s specific involvement in IAV entry via non-clathrin pathways, we examined macropinocytosis and lipid raft-mediated endocytosis. To assess macropinocytosis, we monitored the uptake of 70 kDa Dex, which is specifically internalized via the macropinocytic pathway. While the macropinocytosis inhibitor 5-(N-ethyl-N-isopropyl)-amiloride (EIPA) almost completely inhibited Dex uptake in A549 cells, TRIM62 depletion had no effect (**Fig. 3d**). However, uptake of CTxB, a lipid raft-dependent cargo, was reduced by ∼40% in TRIM62-deficient A549 cells, recapitulating a comparable defect in IAV uptake (**Fig. 3e**). Moreover, we observed colocalization of surface-bound IAV and lipid raft domains at the plasma membrane (as probed by CTxB), as well as colocalization of endocytosed IAV particles with CTxB, suggesting that IAV engages lipid raft pathway during entry, similar to CTxB (**Fig. 3f, g**). Since CTxB can enter via caveolae- and dynamin-dependent or non-caveolar, dynamin-independent lipid raft pathways^33^, we next tested the involvement of caveolae in IAV endocytosis. Intriguingly, Caveolin-1 (CAV1) KD led to an increase in IAV internalization, but pharmacological inhibition of caveolin-1 phosphorylation with genistein had no effect (**Fig. 3h, i**), excluding a role for caveolae in promoting IAV entry. Dynamin inhibition with dynasore also failed to affect IAV uptake, although it strongly impaired CTxB uptake (**Fig. 3i, Supplementary Fig. 4b, c**). By contrast, cholesterol depletion with methyl-β-cyclodextrin (MβCD) severely reduced IAV internalization (**Fig. 3i**). Together, these findings establish that TRIM62 regulates a clathrin-independent, caveolin- and dynamin-independent, but cholesterol-sensitive, lipid raft-mediated entry pathway for IAV.

Since clathrin-independent and lipid raft-dependent pathways of IAV internalization are less studied compared to CME, and several non-clathrin pathways rely on branched F-actin dynamics, we next tested whether branched actin networks are required for driving viral uptake. To address this, we perturbed two key classes of branched actin regulators: the actin-nucleating Arp2/3 complex and its NPFs (**Supplementary Fig. 4d**). Disruption of the Arp2/3 complex, either by siRNA targeting the Arp3 (ACTR3) subunit or by pharmacological inhibition with CK-666, had no effect on IAV uptake (**Fig. 3j**). Similarly, we did not find the involvement of ACTR3 in the uptake of CTxB (**Supplementary Fig. 4e**), suggesting that both CTxB and IAV internalization in A549 cells are branched-actin independent. However, examination of NPFs revealed that while depletion of the surface regulators WAVE2 or N-WASP caused only mild effects, depletion of the endosomal NPF WASH1 markedly impaired IAV internalization (**Fig. 3k**), highlighting an expected role of endosomal WASH1, a subunit of the pentameric WASH complex, in viral entry.

The WASH complex comprises the NPF WASH1 and the SHRC which includes CCDC53, SWIP, Strumpellin, and FAM21. The WASH complex is recruited to endosomes via the retromer subunit VPS35 (**Fig. 3l**). To determine whether all WASH subunits contribute to the regulation of the lipid raft pathway, we individually depleted each subunit, as well as the retromer component VPS35. KD efficiency of the siRNAs was confirmed by RT-PCR (**Supplementary Fig. 4f**). CTxB uptake was then monitored in cells depleted of WASH subunits or VPS35 at 10 and 30 min post-internalization, assessing colocalization with the early endosomal marker EEA1 and Golgi marker GM130, respectively. We found that while WASH1 is a critical regulator of CTxB uptake, SHRC is dispensable (**Fig. 3m, n**). Strikingly, VPS35 depletion enhanced CTxB uptake, in contrast to the effect of WASH1 depletion (**Fig. 3m, n**). Intrigued by these findings, we were curious to examine the distribution of WASH1 and VPS35 relative to lipid rafts. Interestingly, we observed pronounced colocalization of WASH1 and VPS35 with CTxB at the cell surface, an unexpected location given that these proteins are normally localized to endosomes (**Fig. 3o, p**). We also found the colocalization of WASH1 and VPS35 on CTxB-positive vesicles during uptake (**Fig. 3q**), underscoring their involvement in raft-dependent endocytosis. Although WASH1 and VPS35 are typically found associated on endosomal subdomains, we found that their association is cholesterol-dependent. Cholesterol depletion from the plasma membrane using MβCD drastically altered WASH1 distribution and markedly reduced its colocalization with VPS35, as quantified by Manders’ correlation coefficient (**Fig. 3r, s**).

Together, these findings identify WASH1 and VPS35 as reciprocal regulators of lipid raft-mediated endocytosis and implicate TRIM62 in this pathway, which IAV exploits for cell entry.

### WASH subunits and VPS35 distinctly regulate IAV entry

The functional intersection between TRIM62 and the endosomal cargo sorting machinery including WASH and VPS35 in lipid raft pathway prompted us to examine their potential involvement in IAV infection. Surprisingly, an RNAi screen using two independent siRNAs targeting each gene revealed the involvement of VPS35 and every component of WASH in IAV infection. Within the WASH complex, we found striking functional dichotomy: depletion of WASH1, CCDC53, SWIP and Strumpellin impaired IAV infection, whereas FAM21 exhibited an opposite effect. VPS35 depletion similarly enhanced infection, consistent with its functional relationship with FAM21 (**Supplementary Fig. 5a**), suggesting an antiviral FAM21-VPS35 axis.

We next examined how silencing individual WASH subunits or VPS35 affects the protein levels of the other subunits in A549 cells. Western blot analysis showed that depletion of WASH1 or other SHRC subunits destabilized the remaining components, consistent with their function as an interdependent complex. In contrast, VPS35 depletion did not alter the levels of WASH1 or SHRC subunit (CCDC53, FAM21 and SWIP), indicating that VPS35 regulates IAV infection independently of WASH complex integrity (**Supplementary Fig. 5b**). These findings suggest that while WASH1 and the SHRC are directly required for IAV infection, VPS35 may act through its role in WASH recruitment rather than complex stability.

After confirming the involvement of WASH and VPS35 in IAV infection, we next investigated whether these factors contribute to viral entry. To this end, we performed IIF-based assays monitoring sequential steps of IAV entry, beginning with viral binding to the cell surface, followed by internalization and endosomal trafficking. While virus binding was unaffected, IAV entry was altered at downstream steps in cells deficient of WASH subunits and VPS35, with the most pronounced effect observed in M1 uncoating. Consistent with infection phenotypes, depletion of the WASH subunits (WASH1, CCDC53, SWIP, and Strumpellin), blocked viral entry, whereas suppression of FAM21 and the retromer subunit VPS35 showed an opposite phenotype. Along with these factors, TRIM62 was also evaluated, which expectedly showed proviral activity (**Fig. 4a-d**). To gain clearer insight into each factor’s role in viral entry, we plotted average values for each entry step as line graphs. This visualization further emphasized a striking dichotomy: while VPS35 and FAM21 depletion progressively boosted IAV entry, depletion of TRIM62, WASH1 and other SHRC subunits (CCDC53, SWIP, and Strumpellin) led to a gradual decline (**Fig. 4e**). All line graphs pointed to a prominent defect at the uncoating stage. Quantification of cytosolic M1 intensity at the single-cell level confirmed this perturbation and supported the population-level trends (**Fig. 4f**). To further confirm the roles of some key factors, we performed M1 uncoating assay in cells overexpressing them. Overexpression experiments further validated their pro- or antiviral roles in IAV entry: TRIM62-His and GFP-WASH1 significantly boosted uncoating, whereas FAM21-GFP and VPS35-GFP markedly suppressed it (**Fig. 4g, h**). IIF imaging also illustrated a stark divergence in uncoating (M1 dispersal) phenotypes between FAM21/VPS35 depletion and depletion of TRIM62 and other WASH subunits (**Fig. 4i**). Comparing IAV and CTxB uptake, we observed that while SHRC function was essential for IAV internalization, it was dispensable for CTxB uptake, highlighting differences in the lipid raft-mediated uptake of these two cargoes (**Supplementary Fig. 5c**).

**Fig. 4.**
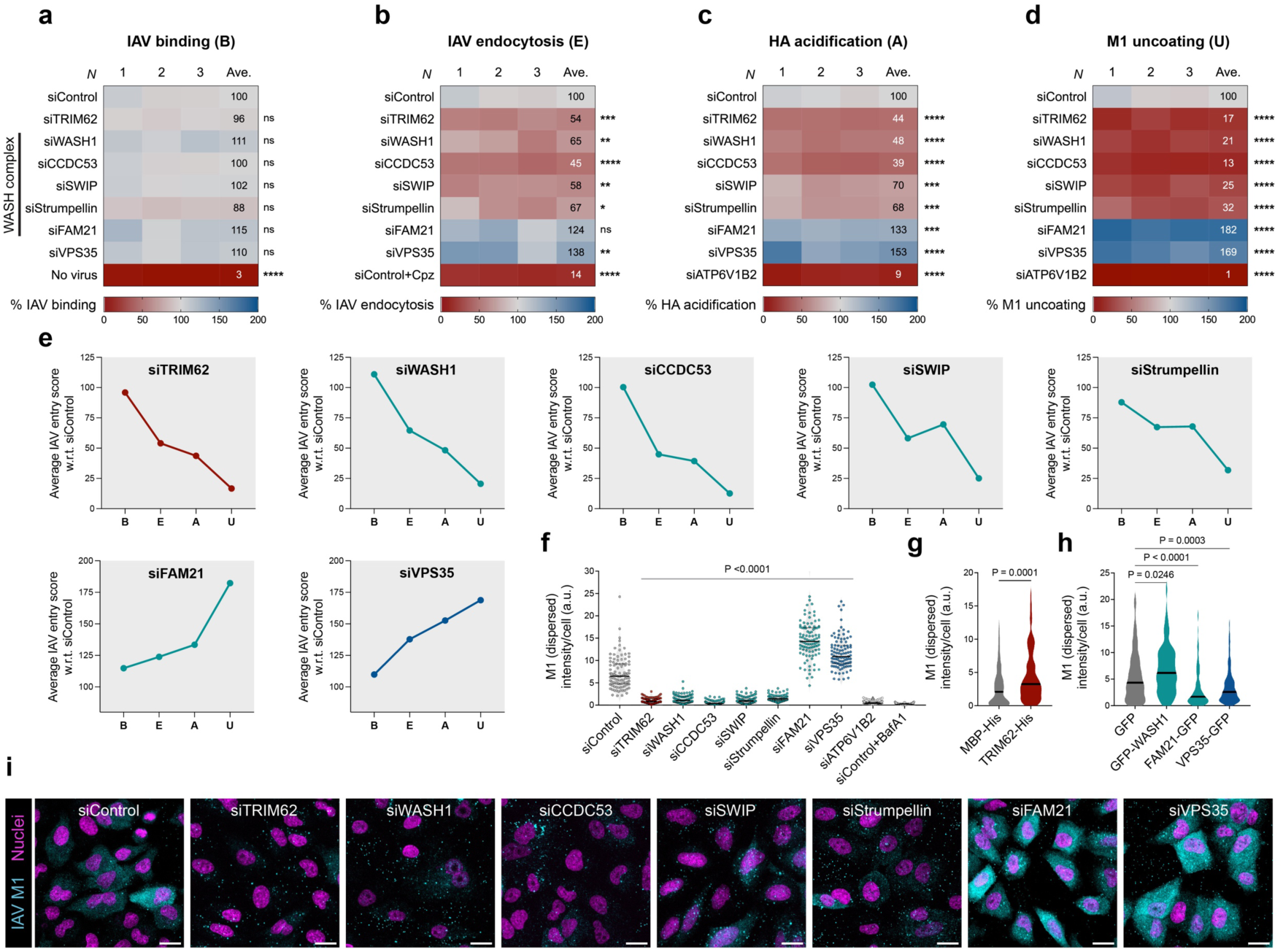
WASH subunits and VPS35 opposingly regulate IAV entry. **a**-**d** Heatmaps showing IAV binding (**a**), IAV endocytosis (**b**), HA acidification (**c**), and M1 uncoating (**d**) in siRNA screens targeting WASH complex subunits and VPS35. All data are representative of three biological replicates. Average values of three replicates are shown. *n* > 500 cells per sample. **e** Line graphs showing average values for IAV binding (B), endocytosis (E), HA acidification (A), and M1 uncoating (U), upon KD of individual factors. **f** Scatter plot showing the quantification of cytosolic M1 (dispersed) intensity per cell. *n* ≥ 99 cells per sample. **g** Quantification of cytosolic M1 (dispersed) intensity in cells expressing MBP-His or TRIM62-His at 120 min post-infection. *n* = 120 cells per sample. **h** Quantification of cytosolic M1 (dispersed) intensity in cells expressing GFP and GFP-tagged WASH1, FAM21 or VPS35 at 120 min post-infection. *n* ≥ 125 cells per sample. **i** High-content confocal images showing M1 uncoating (M1, cyan and nuclei, magenta) in KD cells. Data in **a**-**d**,**f,h** were analyzed using one-way ANOVA using multiple-comparisons. Data in **g** was analyzed using unpaired two-tailed *t*-test, P values are indicated. P values < 0.05 were considered significant.

Collectively, our findings emphasize a crucial role of WASH and VPS35 in differentially regulating IAV entry. While WASH1, CCDC53, SWIP and Strumpellin promote viral entry, the FAM21-VPS35 axis plays an antiviral role. Importantly, the effects of TRIM62 depletion closely resembled those of WASH1 and other SHRC subunits except FAM21 across viral entry steps, suggesting that TRIM62 may act in concert with the WASH complex to regulate this noncanonical endocytic pathway.

### TRIM62 interacts with the WASH complex through its C-terminus, regulating WASH-VPS35 association and IAV entry

The similar phenotypes observed upon TRIM62 and WASH1 depletion on IAV internalization raised the possibility that these factors functionally cooperate to regulate lipid raft-mediated IAV entry. To explore this, we first examined whether TRIM62 and WASH1 colocalize at sites of viral uptake. Airyscan high-resolution microscopy in A549 cells revealed an enrichment of TRIM62-Myc in branched actin-rich lamellipodia, where endogenous WASH1 was also detected (**Fig. 5a, b**). Although the lamellipodial presence of the WASH complex has been debated, as it is generally considered endosome-associated ^24,34^, our observations in A549 cells clearly demonstrate that WASH1 localizes to the lamellipodium, colocalizing with TRIM62. Extending these observations, we also detected colocalization of TRIM62 with FAM21, further supporting its spatial association with the WASH complex (**Fig. 5c, d**).

**Fig. 5.**
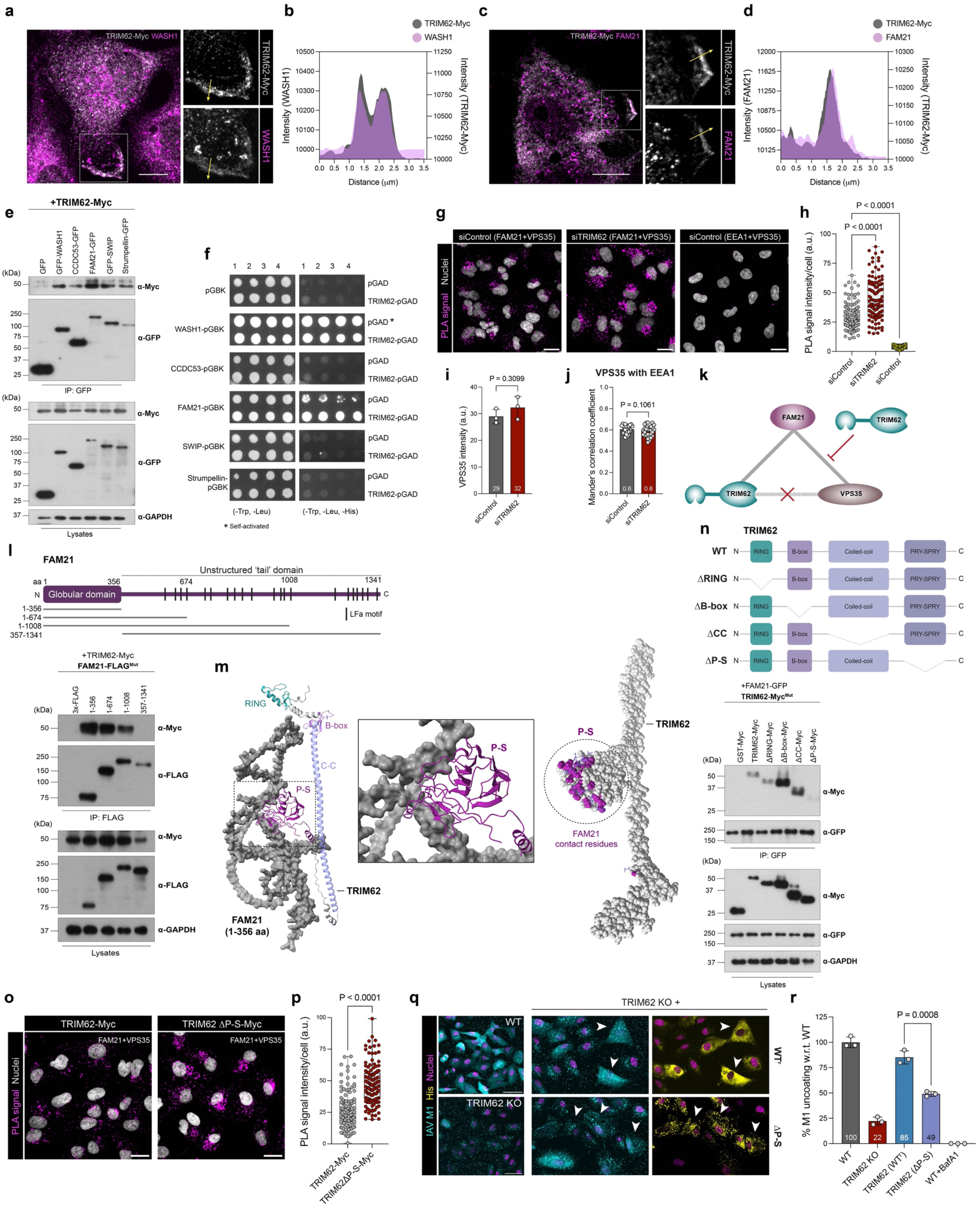
TRIM62 interacts with FAM21 via its C-terminus, regulating WASH-VPS35 association and thereby promoting IAV entry. **a** High-resolution microscopy image of a representative cell showing the presence of endogenous WASH1 (magenta) and overexpressed TRIM62-Myc (grey) at lamellipodium. Insets show colocalization of TRIM62-Myc, and WASH1 at the lamellipodium. Scale bars, 10 µm. **b** Fluorescence intensity profiles of TRIM62-myc and WASH1 along the yellow line are shown. **c** High-resolution microscopy image of a representative cell showing the presence of endogenous FAM21 (magenta) and overexpressed TRIM62-Myc (grey) at lamellipodium. Insets show colocalization of TRIM62-Myc, and FAM21 at lamellipodium. Scale bars, 10 µm. **d** Fluorescence intensity profiles of TRIM62-myc and FAM21 along the yellow line are shown. **e** Pull-down assay showing interactions between TRIM62 and WASH subunits. HEK293T cells were co-transfected with TRIM62-Myc and GFP-tagged constructs of WASH subunits including WASH1, CCDC53, SWIP, Strumpellin and FAM21, and lysates were incubated with GFP-Trap beads. Western blotting was done to detect TRIM62-Myc in the precipitates. **f** Y2H assay showing a direct interaction between TRIM62 and FAM21. Yeast co-transformants were spotted on -Trp, -Leu plates (left) to indicate growth, and -Trp, -Leu, -His plates (right) to confirm interaction. * Indicates self-activation for WASH1. **g**,**h** High-content confocal images (**g**) and quantification (**h**) of PLA signals from FAM21-VPS35, and EEA1-VPS35 interactions. Scale bars, 20 µm. *n* = 118 cells per sample. **i** Quantification of VPS35 intensity. *n* = 1029 cells (siControl), *n* = 839 cells (siTRIM62). **j** Quantification of VPS35-EEA1 colocalization based on Mander’s correlation coefficient. *n* > 1800 cells per sample. **k** Schematic showing FAM21 interactions with TRIM62 and VPS35, with TRIM62 negatively regulating the FAM21-VPS35 interaction. **l** Schematic showing FAM21 domain organization (top). Pull-down assay (bottom) showing interactions between TRIM62 and FAM21 truncation mutants. HEK293T cells were co-transfected with TRIM62-Myc and FAM21-FLAG mutant constructs, and lysates were incubated with FLAG-immunotrap beads. Western blotting was done to detect TRIM62-Myc in the precipitates. **m** AlphaFold 3 predictions showing interaction between FAM21 N-terminal globular region (1-356 aa) and TRIM62 and the contact residues are highlighted (magenta). **n** Schematic showing TRIM62 domain organization (top). Pull-down assay (bottom) showing interactions between FAM21 and TRIM62 domain-deleted mutants. HEK293T cells were co-transfected with FAM21-GFP and TRIM62-Myc mutant constructs, and lysates were incubated with GFP-Trap beads. Western blotting was done to detect TRIM62-Myc in the precipitates. **o,p** High-content confocal images (**o**) and quantification (**p**) of PLA signals from VPS35-FAM21 interaction in cells stably expressing either TRIM62-Myc or TRIM62ΔP-S-Myc. Scale bars, 20 µm. *n* = 130 cells per sample. **q** High-content confocal images showing M1 uncoating in WT, TRIM62 KO, and KO cells stably expressing WT TRIM62-His or its mutants. M1 (cyan), His (yellow), and nuclei (magenta). Scale bars, 50 µm. **r** Quantification of % M1 uncoating, normalized to WT. *n* > 450 cells per sample. Data in **h,r** were analyzed using one-way ANOVA using multiple-comparisons. Data in **i,j,p** was analyzed using unpaired two-tailed *t*-test, P values are indicated. P values < 0.05 were considered significant. All bar graphs show the mean of n = 3 ± SD.

We next investigated whether TRIM62 physically interacts with the WASH complex. We performed pulldown assays using GFP-nanotrap immunocapture on extracts from HEK293T cells that co-expressed TRIM62-Myc and GFP-tagged components of the WASH complex, and found that TRIM62-Myc interacted with all WASH subunits (**Fig. 5e**). However,

no interaction was detected with VPS35 or other retromer subunits (**Supplementary Fig. 6a**), suggesting TRIM62’s specific association with WASH. Next, we examined TRIM62’s direct interaction with WASH subunits by Y2H assay. Since yeast lacks conserved WASH components^35,36^, this system was chosen to detect direct interactions without involvement of endogenous partners. Among the WASH subunits, yeast growth was observed in the presence of FAM21, despite some degree of self-activation, indicating a direct interaction with TRIM62. However, strong self-activation WASH1 in yeast precluded reliable determination of its direct interaction with TRIM62 (**Fig. 5f**).

Since TRIM62 interacts with the WASH complex via FAM21 but does not bind retromer, we asked whether this interaction influences WASH-retromer association. To address this, we examined WASH recruitment to retromer-enriched foci by assessing FAM21-VPS35 interaction using proximity ligation assay (PLA). TRIM62-deficient A549 cells showed significantly enhanced PLA signals, despite unchanged VPS35 levels (**Fig. 5g-i**). As a negative control, we checked PLA signals between EEA1 and VPS35, proteins that are not known to interact despite their endosomal presence. Negligible PLA signals from EEA1-VPS35 interaction confirmed assay specificity (**Fig. 5g, h**). VPS35 association with early endosomes remain unchanged (**Fig. 5j**), suggesting that the enhanced PLA signals from FAM21-VPS35 interaction were not due to an increased presence of VPS35 on early endosomes. To examine whether FAM21 alone or the WASH complex as a whole associates more strongly with VPS35, we performed PLA between WASH1 and VPS35 in TRIM62-deficient A549 cells. The enhanced WASH1-VPS35 PLA signals indicated increased recruitment of the WASH complex to VPS35 (**Supplementary Fig. 6b, c**). We next addressed whether the enhanced association of the WASH complex with VPS35 is specific to the A549 cell line. To this end, we checked FAM21-VPS35 interaction in TRIM62-deficient HeLa cells and found markedly elevated PLA signals confirming that the effect is general and not cell type specific (**Supplementary Fig. 6d, e**). Together, these data suggest TRIM62’s regulatory role in the association between the WASH complex and VPS35 through limiting FAM21-VPS35 interaction (**Fig. 5k**).

We next mapped the domains responsible for TRIM62-FAM21 interaction. Using truncation mutants we found that the N-terminal globular region (1-356 amino acids) of FAM21, which includes the “head” domain (1-220 amino acids), mediates the interaction with TRIM62 (**Fig. 5l**). This “head” domain of FAM21 is essential for its incorporation into the WASH complex^37–39^, while its ∼1100 amino acid-long disordered “tail” region, which harbours multiple leucine-phenylalanine-acidic (LFa) motifs, mediates endosomal anchoring via interaction with VPS35^38,40,41^. We employed AlphaFold 3.0 modelling, which predicted an interaction interface between the FAM21 globular region and the P-S domain of TRIM62 (**Fig. 5m**). Consistent with this prediction, co-immunoprecipitation assays with TRIM62 deletion mutants revealed that the N-terminal globular region of FAM21 interacts specifically with the C-terminal P-S domain of TRIM62. Loss of the P-S domain abolished FAM21 binding, while deletions of the RING, B-box, or CC domains did not (**Fig. 5n**). We also found that the P-S domain of TRIM62, along with its CC domain, is critical for its interaction with WASH1 (**Supplementary Fig. 6f**). To confirm whether TRIM62’s interaction with FAM21 via the P-S domain is critical for regulating retromer-WASH association, we generated A549 cells stably expressing either TRIM62-Myc or a P-S domain truncation mutant (TRIM62ΔP-S-Myc), and examined VPS35-FAM21 association by PLA. Cells expressing TRIM62ΔP-S-Myc displayed significantly enhanced PLA signals relative to TRIM62-Myc (**Fig. 5o, p**), indicating that the P-S domain negatively regulates the FAM21-VPS35 interaction, thereby modulating retromer-WASH association.

To delineate the functional relevance of this interaction in IAV entry, we performed rescue experiments in KO cells. Stable expression of WT TRIM62 (WT^r^) in the KO background, almost fully rescued the M1 uncoating defect observed in KO cells, whereas the P-S domain-truncated mutant (ΔP-S) failed to rescue it (**Fig. 5q, r**). Interestingly, we found that while the RING mutant lacking E3 ubiquitin ligase activity (RING^m^) or the RING domain deletion mutant (ΔRING) fully rescued uncoating in KO cells, mutants lacking the B-box (ΔB-box) or CC (ΔCC) domains failed to do so (**Supplementary Fig. 6g**).

These findings identify the C-terminal CC and P-S domains of TRIM62 as critical for its interaction with the WASH complex and for regulating IAV entry, establishing TRIM62 as a key modulator of WASH-retromer dynamics during viral internalization and intracellular trafficking.

### TRIM62 and VPS35 antagonistically regulate WASH endosome-cytosol distribution with distinct effects on IAV entry

The increased FAM21-VPS35 interaction observed in TRIM62-deficient cells suggested that TRIM62 modulates the spatial distribution of the WASH complex within the endosomal network. To address this, we examined WASH1 and FAM21 distribution upon TRIM62 suppression. In TRIM62 KD cells, we observed enhanced colocalization of both WASH1 and FAM21 with EEA1-positive early endosomes compared to controls (**Fig. 6a–d**), along with increased WASH1 and FAM21 intensity on endosomes (**Supplementary Fig. 7a, b**). In addition to early endosomes, we also examined the presence of WASH1 on Rab7^+^ endosomes in TRIM62 KD cells by high-resolution live-cell imaging. We co-expressed green fluorescent protein (GFP)-tagged WASH1 (GFP-WASH1) and red fluorescent protein (RFP)-tagged Rab7 (RFP-Rab7) in A549 cells and treated them with the PIKfyve inhibitor apilimod to induce endosomal enlargement^42^, thereby enabling clearer visualization of WASH subdomains on endosomes. We observed that compared to controls, TRIM62 depletion markedly enhanced WASH1 recruitment to Rab7^+^ endocytic vesicles (**Fig. 6e**). We next asked whether WASH enrichment on endosomes was driven by protein abundance, and therefore, checked the levels of individual WASH subunits and VPS35 in TRIM62 KD or KO cells. However, we observed no appreciable differences in the protein levels (**Fig. 6f, Supplementary Fig. 7c**), indicating that TRIM62 regulates WASH subcellular distribution without altering its overall abundance.

**Fig. 6.**
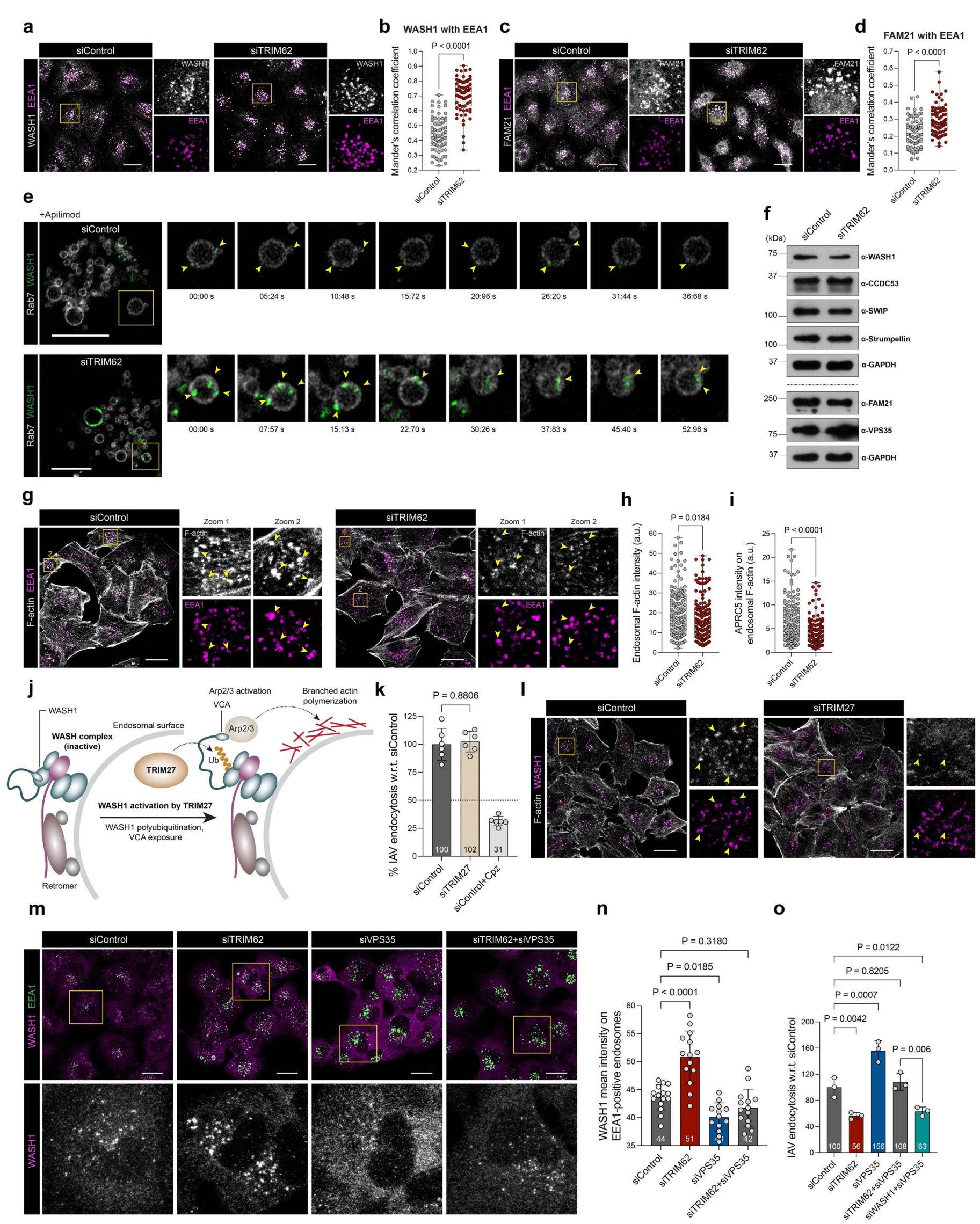
WASH1 mediates IAV entry independent of its F-actin nucleation activity, while TRIM62 counteracts VPS35 to promote WASH1-driven viral entry. **a** Confocal microscopy images showing WASH1 (grey) and EEA1-positive endosomes (magenta) in control and TRIM62-depleted cells. Insets show the distribution of WASH1 and EEA1. Scale bars, 20 µm. **b** Quantification of WASH1-EEA1 colocalization based on Mander’s correlation coefficient. *n* = 70 cells per sample. **c** Confocal microscopy images showing FAM21 (grey) and EEA1-positive endosomes (magenta) in control and TRIM62-depleted cells. Insets show the distribution of FAM21 and EEA1. Scale bars, 20 µm. **d** Quantification of FAM21-EEA1 colocalization based on Mander’s correlation coefficient. *n* = 74 cells per sample. **e** High-resolution, time-lapse images of a representative endosome (outlined by the box) in a cell expressing RFP-Rab7 (grey) and GFP-WASH1 (green) upon apilimod (1 µM) treatment in control and TRIM62 KD cell. Foci of GFP-WASH1 on the endosome are indicated by yellow arrowheads. Scale bars, 10 µm. **f** Western blots showing the levels of WASH complex proteins and VPS35 in control and TRIM62 KD cells. **g** Confocal microscopy images showing F-actin (grey) and EEA1-positive endosomes (magenta) in control and TRIM62 KD cells. Yellow arrowheads within the insets show the distribution of F-actin at endosomes. Scale bar, 20 µm. **h** Scatter plot showing F-actin intensity on EEA1+ endosomes. *n* = 150 F-actin patches per sample. **i** Scatter plot showing ARPC5 intensity in F-actin area on Rab5QL+ endosomes. *n* = 150 F-actin patches per sample. Related to Supplementary Fig. 7d. **j** Schematic showing WASH1 activation by TRIM27. TRIM27 ubiquitinates WASH1 exposing the VCA domain which is crucial for the recruitment of Arp2/3 for branched actin nucleation at endosomes. **k** Quantification of percentage of IAV endocytosis at 30-min post-infection, in control and TRIM27 KD cells. *n* = 4000 cells per sample. **l** Confocal microscopy images showing F-actin (grey) and WASH1-positive endosomes (magenta) in control and TRIM27-depleted cells. Yellow arrowheads within the insets show the distribution of F-actin at endosomes. Scale bar, 20 µm. **m** Confocal images showing WASH1 (magenta) and EEA1 (green) in control and TRIM62, VPS35 and TRIM62+VPS35 KD cells. Lower panel shows zoomed-in images of WASH1. Scale bars, 20 µm. **n** Quantification of WASH1 intensity on EEA1+ endosomes. *n* > 70 cells per sample. **o** Quantification of percentage of IAV endocytosis. *n* > 1500 cells per sample. Data in **k,n,o** were analyzed using one-way ANOVA using multiple-comparisons. Data in **b,d,h,i** were analyzed using unpaired two-tailed *t*-test, P values are indicated. P values < 0.05 were considered significant. All bar graphs show the mean of n = 3 ± SD.

Observing enhanced localization of WASH on endosomes, we were prompted to examine whether this was accompanied by increased branched F-actin assembly at endosomes driven by WASH’s NPF activity. Surprisingly, we found that F-actin signals coinciding EEA1-positive early endosomes were significantly reduced in TRIM62-deficient cells compared to controls (**Fig. 6g, h**). Since WASH promotes F-actin assembly by activating the Arp2/3 complex, we next examined ARPC5, an Arp2/3 subunit, in cells overexpressing GFP-Rab5Q67L, a constitutively active GTP-bound form of Rab5 that induces endosomal enlargement. ARPC5 intensity was measured on F-actin patches and found to be reduced in TRIM62-deficient cells compared to controls (**Fig. 6i, Supplementary Fig. 7d**). These findings indicate that despite enhanced recruitment of WASH to endosomes in the absence of TRIM62, its NPF function remains impaired.

We next asked whether this impaired WASH activity affected endosomal trafficking of prototypical cargoes of retromer/WASH such as glucose transporter 1 (GLUT1), and epidermal growth factor receptor (EGFR). VPS35 and FAM21 depletion mislocalize endogenous GLUT1 from the cell surface to late endosomes/lysosomes and the Golgi, respectively^43,44^. Interestingly, TRIM62 depletion in HeLa cells caused steady-state mislocalization of GLUT1, which accumulated in the Golgi as verified by brefeldin A- and monensin-induced Golgi perturbation (**Supplementary Fig. 8a, b**). Additionally, TRIM62 deficiency in A549 cells led to an increased presence of EGFR in LAMP1-positive vesicles at steady state compared to controls (**Supplementary Fig. 8c, d**). Together, these findings uncover a previously unrecognized role for TRIM62 in maintaining endosomal cargo trafficking.

Although we found that global Arp2/3 inhibition does not interfere with viral entry (**Fig. 3j**), but TRIM62 deficiency renders WASH inactive for branched actin nucleation, we addressed whether viral entry defects result specifically from the loss of WASH’s NPF activity. To test this, we downregulated TRIM27, another TRIM protein known to specifically activate WASH1 through K63-linked ubiquitination, thereby exposing its VCA domain, which then stimulates the Arp2/3 complex to drive actin nucleation at endosomes (**Fig. 6j**). Interestingly, depletion of TRIM27 using a previously validated siRNA^45^ did not cause any reduction in IAV endocytosis (**Fig. 6k**), but led to a marked reduction in endosomal F-actin assembly (**Fig. 6l**) Thus, while TRIM62 depletion impairs canonical, NPF-dependent cargo trafficking activity of WASH1, the block in viral entry reveals a distinct, noncanonical, actin-independent role of WASH1.

Finally, we examined how depletion of TRIM62 or VPS35, alone or in combination, alters WASH1 distribution and IAV entry. Since TRIM62 restricts WASH endosomal localization, displaying a functional antagonism with VPS35, we probed the functional interplay of these factors during IAV entry. As expected, TRIM62 depletion enhanced WASH1 localization on endosomes and impaired IAV entry, whereas VPS35 loss caused WASH1 to disperse into the cytosol and promoted viral entry. Remarkably, co-depletion of TRIM62 and VPS35 normalized WASH1 endosomal distribution and restored viral endocytosis to near-control levels (**Fig. 6m-o**). In co-depleted cells where IAV internalization was restored, endosomal actin remained reduced (**Supplementary Fig. 8e**), further confirming that WASH1 supports IAV entry independently of its NPF activity. Simultaneous depletion of VPS35 and WASH1, however, significantly impaired IAV uptake relative to TRIM62-VPS35 co-depletion, underscoring WASH1’s essential role in viral endocytosis (**Fig. 6o**).

Together, these findings establish TRIM62 as a key antagonist of VPS35 in regulating WASH complex recruitment to endosomes, maintaining WASH1’s cytosolic presence to facilitate productive IAV entry.

## Discussion

The intricately coordinated entry program of IAV relies on a dynamic and finely balanced endosomal network, which helps the virus internalize and move deep into the cytoplasm, bypassing cortical barriers and cytosolic crowding^46,47^. This study identifies TRIM62 as a mediator of IAV endocytosis through a clathrin-independent, lipid raft-mediated pathway and downstream trafficking, and implicates the WASH complex and retromer component VPS35 in this process. Importantly, we find that TRIM62’s canonical E3 ubiquitin ligase activity is dispensable for viral entry, highlighting its scaffolding function through the other domains. Our data further reveal that TRIM62 interacts with the WASH complex via its FAM21 subunit and that the WASH complex contributes to IAV entry. Notably, we observed a striking functional dichotomy within the WASH complex: WASH1, CCDC53, SWIP, and Strumpellin promote viral entry, whereas FAM21 exerts an opposing effect. FAM21’s functional divergence from the otherwise supportive roles of other WASH complex subunits in IAV entry implies a potentially unique regulatory mechanism that remains to be elucidated. Notably, previous studies also reported WASH-independent role of FAM21, highlighting its functional versatility and distinguishing it from other WASH subunits^48,49^.

A recent preprint described a clathrin-independent endocytic pathway for human papillomavirus (HPV), driven by WASH1 through its NPF activity^23^. In this study, the authors found WASH1’s involvement in the late stage of viral endocytosis, where it promotes endocytic vesicle scission via actin polymerization. Although both this study and ours highlight a critical role for WASH1 in HPV and IAV endocytosis, respectively, the SHRC subunits were found to exhibit divergent functions. Contrasting to our findings on IAV entry, CCDC53, SWIP, and Strumpellin showed no role in HPV entry, while FAM21 had only a modest effect^23^. Given the distinct trafficking routes of IAV and HPV, where IAV follows the degradative pathway to reach late endosomes and HPV traffics to the *trans*-Golgi, we reason that the entry requirements of these two viruses are different. Another key difference between WASH1’s role in HPV and IAV entry lies in its NPF activity: while HPV relies on WASH1-mediated branched actin nucleation at the plasma membrane for its endocytosis, IAV endocytosis in A549 cells does not require branched actin, but still depends on WASH1. The non-essentiality of WASH1’s NPF activity in IAV entry is supported by the observation that depletion of TRIM27 does not impair viral endocytosis. Consistent with previous findings that TRIM27’s absence fails to activate WASH1 for its NPF activity^45^, we also observed a marked reduction in endosomal F-actin upon TRIM27 depletion, yet viral internalization remained unaffected. These findings highlight a noncanonical, actin-independent role for WASH1 in IAV endocytosis, likely through its scaffolding function. However, the precise mechanism by which WASH1 drives viral internalization remains to be elucidated.

While examining lipid raft-mediated endocytosis using the model cargo and lipid raft marker CTxB, we unexpectedly found the presence on WASH1 and VPS35 at the plasma membrane, colocalizing with CTxB. Importantly, both WASH1 and VPS35 exert opposite functions in the cellular uptake of CTxB and IAV, pointing at their common regulatory role. Presence of IAV particles in lipid rafts at the plasma membrane and in endosomes indicates viral entry through a lipid raft-mediated pathway, where WASH1 and VPS35 operate antagonistically. Notably, a previous study investigating vaccinia virus (VV) entry found the WASH subunit FAM21 to colocalize with the plasma membrane lipid raft marker CTxB and VV in HeLa cells, suggesting that WASH associates with the lipid raft microdomains on the plasma membrane^50^, similar to our observations. The same group later reported that WASH1 and FAM21 play critical roles in VV entry in HeLa cells, whereas the other WASH subunits are dispensable^51^. Our findings on IAV, together with previous studies on HPV and VV, indicate that individual WASH subunits differentially regulate endocytic processes depending on the virus.

Our data suggest that TRIM62 opposes VPS35 function in WASH recruitment to endosomes. By directly interacting with FAM21, TRIM62 limits WASH’s endosomal recruitment and likely increases its availability at the plasma membrane to drive IAV endocytosis (**Fig. 7**). The importance of WASH’s cytosolic presence in viral entry is evident in VPS35-deficient cells, where WASH redistributes from endosomes to the cytosol, increasing its cytosolic abundance and thereby enhancing viral endocytosis efficiency. As WASH’s endosome-cytosol balance is restored in cells co-depleted of TRIM62 and VPS35, viral endocytosis normalizes to basal levels, underscoring the importance of the delicate balancing act between these two proteins in regulating WASH distribution for IAV endocytosis. Notably, despite restoration of IAV entry in co-depleted cells, endosomal actin remained reduced, as observed in cells depleted of either TRIM62 or VPS35, indicating that the WASH’s NPF activity is not a determinant of IAV entry, but its spatial distribution is critical.

**Fig. 7.**
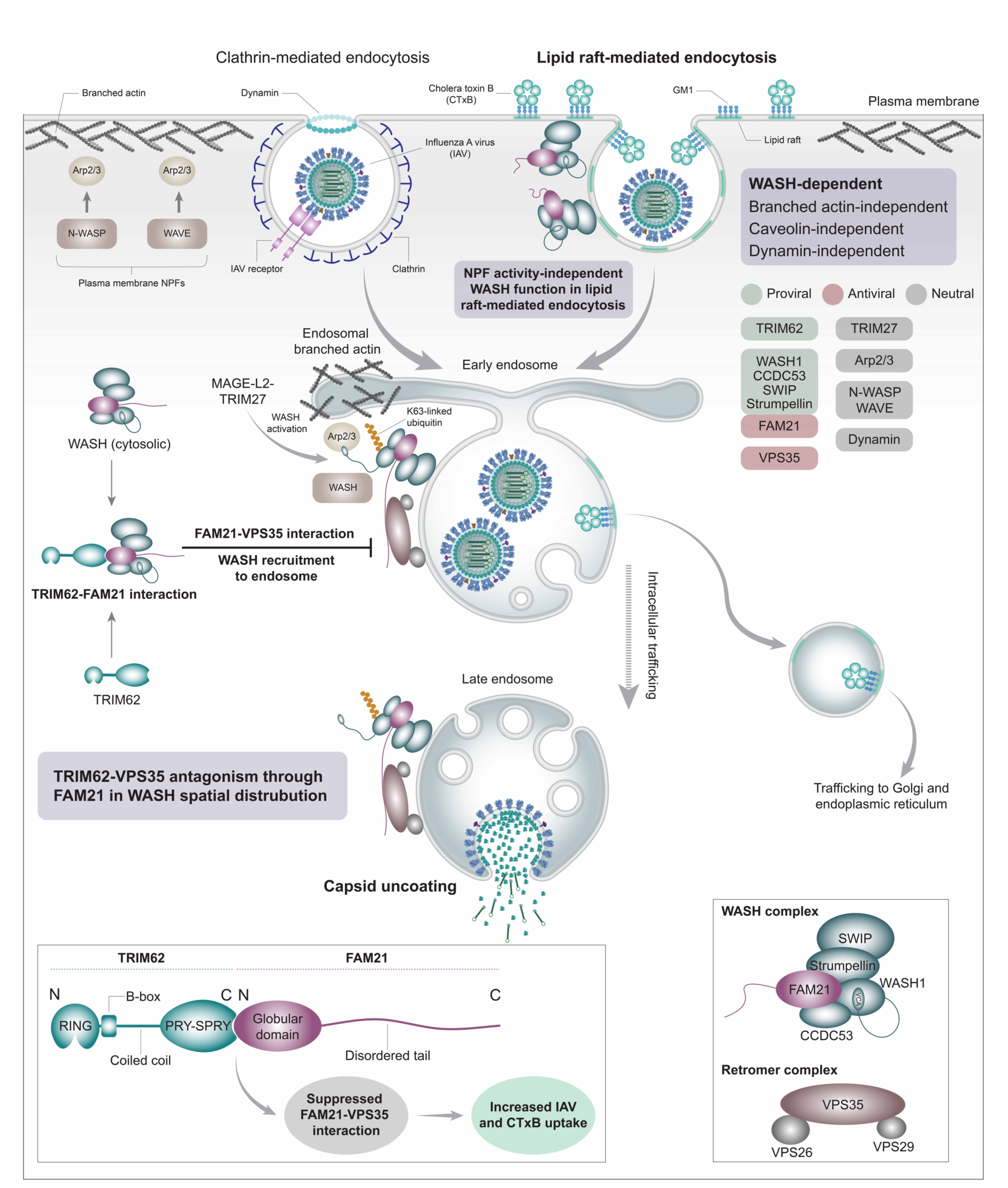
Model showing opposing regulation of the WASH complex by TRIM62 and VPS35 during IAV entry. IAV enters host cells via the classical clathrin-mediated endocytosis (CME) and a WASH-dependent, lipid raft-mediated pathway. This WASH-dependent pathway is shared by IAV and CTxB, but with different endocytic requirements. While IAV requires WASH for its entry through this lipid raft pathway, its internalization is not dependent on branched actin, caveolin, or dynamin. Both CME and lipid raft pathways converge at early endosomes, where further trafficking and cargo sorting occur. In the cytosol, TRIM62 directly interacts with FAM21 of the WASH complex via its C-terminal PRY-SPRY domain, and inhibits WASH’s recruitment to endosomes by limiting FAM21-VPS35 interaction. Thus, TRIM62 and VPS35 oppose each other in modulating WASH distribution and differentially regulating IAV entry.

Taken together, our study reveals a TRIM62-WASH-VPS35 axis in the regulation of IAV entry and establishes an NPF activity-independent paradigm in WASH biology. WASH possibly serves as a scaffolding hub that coordinates multiple factors involved in endocytosis and membrane trafficking. Mechanistically, while our study highlights TRIM62’s pivotal role in regulating WASH dynamics through limiting FAM21-VPS35 interaction, thereby promoting IAV entry, further studies are required to gain clearer insights into WASH’s precise role in viral endocytosis.

## Methods

### Cell culture

A549, MDCK, HeLa, HepG2, and HEK293T cells were purchased from American Type Culture Collection (ATCC) and Huh-7 was purchased from Merck, and cultured in DMEM (Gibco, 10566016), supplemented with 10% fetal bovine serum (FBS) (Merck, F2442), 1x MEM non-essential amino acids (Gibco, 11140050), 1x penicillin-streptomycin-glutamine (Gibco, 10378016) and were maintained at 37 °C in 5% CO_2_. Cells were maintained at 70-80% confluency. TRIM62 knockout (KO) cells were maintained at 60% confluency.

### Virus, bacterial, and yeast strains

Purified IAV X-31 (H3N2) was purchased from Microbiologics (previously Virapur), USA and stored at -80 °C. The median tissue culture infectious dose (TCID50) of purified X-31 virus was determined as 1.95×10^9^ infectious units/ml. The IAV strains NYMC_X311 (A/Brisbane/1/2018) (H3N2), Udorn (A/Udorn/72) (H3N2), and WSN (A/WSN/1933) (H1N1) were kind gifts from Yohei Yamauchi (ETH Zurich). DENV-2 was a kind gift from Arup Banerjee (RCB, Faridabad). All strains were propagated in MDCK cells and stored at -80 °C. All the bacterial transformations were done in chemically competent Stbl3 strain of *E. coli*. Yeast 2-hybrid experiments were performed in the Y2HGold strain of *Saccharomyces cerevisiae* (Takara Bio, Inc.) (kind gift from Mahak Sharma, IISER Mohali).

### Plasmids and transfection

All the plasmids used in this study are described in the Supplementary Table 1. Plasmids for transient overexpression were generated by inserting amplified DNA into pEGFP-N1 or pEGFP-C1. Plasmids for stable cell line generation were generated by inserting amplified DNA into pLenti CMV GFP puro (Addgene, 17448). For plasmid transfection into A549 cells, Lipofectamine™ LTX Reagent with PLUS™ Reagent (Invitrogen, 15338100) was used as per manufacturer’s protocol. For transfection in HEK293T cells, 1 µg of both plasmids were transfected together using 6 µl of 0.1% polyethylenimine (PEI, Merck, 408727) (1:3 ratio). Just before the plasmid transfection, the cell medium was changed with warm supplemented DMEM.

### Generation of TRIM62 knockout (KO) cell line

TRIM62 was knocked out in A549 cells using the CRISPR/Cas9 approach. Guide RNA (gRNA) sequence (GCAGCTGCTCAAGCGACAAC) targeting TRIM62 genomic region was designed using CRISPOR and was cloned in pSpCas9(BB)-2A-Puro (PX459) vector. This plasmid was transfected into A549 cells using the Cell Line Nucleofector® 2b Kit T (Lonza, VCA-1002) according to the manufacturer’s protocol. In summary, 1 x 10^6^ cells were taken and centrifuged at 90 g for 10 min and the pellet was resuspended in 100 µl Solution T. Immediately, 4 µg plasmid DNA was added to it and the sample was loaded into the cuvette and pulsed twice using programme X-001. Cells were incubated at 37 °C for 3 days and then passaged and confirmed by sequencing and quantitative reverse transcription polymerase chain reaction (RT-PCR).

### Generation of stable cell lines

TRIM62 WT and mutants were stably expressed in TRIM62 KO cells using the lentiviral vector system. To generate lentiviruses, HEK293T cells were transfected with pCMVR8.74 and pMD2.G lentiviral packaging plasmids along with TRIM62-His (wild type, WT) or mutants cloned in the pCMV lenti puro vector, using PEI. 6-8 h after transfection. Cells were washed with 1x phosphate-buffered saline (PBS) and prewarmed media was added. The viruses were collected after 72 h of transfection and centrifuged to clear the supernatant. A549 cells were transduced with fresh lentiviral particles using polybrene (Merck, H9268). After 60-72 h of transduction, cells were passaged and used for experiments after confirming the expression of His using indirect immunofluorescence (IIF).

### siRNA transfection

siRNAs targeting human genes were purchased from Qiagen. AllStars Negative Control (siControl) was used as negative control. siATP6V1B2 was used as a positive control for IAV infection. For siRNA knockdown, A549 cells were reverse transfected with 10 nM siRNA using Lipofectamine™ RNAimax (Invitrogen, 13778-150) and incubated for 72 h. All siRNAs used in this study are listed in Supplementary Table 2. Knockdown efficiency was validated by RT-PCR or western blotting. The primers used for RT-PCR used in this study are given in Supplementary Table 3. RT-PCR was performed using iTaq™ Universal SYBR® Green Supermix (Bio-Rad, 1725121) on a CFX96 system (Bio-Rad). Data analysis was performed using 2-ΔΔCT method.

### RNAi screening of TRIM proteins

Three different siRNA sequences targeting 20 TRIM proteins were purchased from Qiagen. A549 cells were reverse transfected with 10 nM siRNAs using Lipofectamine RNAiMAX and incubated for 72 h at 37 °C in CO_2_ incubator. After incubation, cells were infected with IAV (purified X-31) at an MOI of 0.01 for 10 h. The virus was diluted in infection medium (50 mM HEPES, 0.2% BSA in DMEM). Bafilomycin A1 (BafA1) (Merck, B1793) was used as an inhibitor of virus infection. After 10 h, cells were washed with PBS and fixed with 4% formaldehyde for 20 min at room temperature (RT) and then IIF was performed to detect viral nucleoprotein (NP). Cells were imaged using a high-content confocal quantitative image cytometer (CQ1, Yokogawa). NP intensity was quantified using ImageJ software and average intensity per cell was calculated. 15 TRIMs were selected out of 20 if two or more siRNAs showed more than 50% decrease in NP intensity. These selected 15 TRIM proteins were screened for viral ribonucleoprotein (vRNP) nuclear import.

For vRNP nuclear import screen, cells were transfected with siRNAs and after 72 h, cells were incubated with 0.25 µl of 1.95×10^9^ infectious units/ml of IAV X-31 per well of 96-well plate for 4 h. For nuclear import assay of incoming virus particles, virus was diluted in 1 mM CHX (Merck, C1988) containing infection media. After 2 h, infection was removed and fresh CHX containing media was added for another 2 h. In control wells, importazole (Ipz, 40 nM) (Cell Signalling Technology, 10451s) was also added throughout the assay. After incubation at 37 °C, cells were washed with PBS and fixed with 4% formaldehyde for 20 min at RT and then IIF was performed to stain viral NP. Cells were imaged using CQ1.

The experiments were carried out in triplicate and nine (3×3) z-stack images were acquired from random fields from a single well of a 96-well optical bottom plate (Greiner, 655090). Imaging was performed using a 20x or 40x objective lens in CQ1. The images were stacked and maximum intensity projections were taken for image analysis using ImageJ.

### Indirect immunofluorescence (IIF)

Fixed cells were permeabilized with permeabilization solution (PS) (5% FBS, 1% BSA, and 0.1% saponin in 1x PBS) for 30 min at RT. Cells were incubated with primary antibody diluted in PS for 2 h at RT and then washed three times with PBS for 10 min each. After washing, it was incubated with Alexa Fluor conjugated secondary antibody for 1 h at RT. Nucleus (DNA) was stained with Hoechst (Thermo Scientific, 62249). The plate was washed three times, 10 min each with PBS. Imaging was performed in CQ1. Image analysis was performed using CellProfiler, KNIME, and ImageJ software.

Primary antibodies used were mouse anti IAV-NP (HB65, 1:50), mouse anti IAV-M1 (HB64, 1:30) from ATCC; mouse anti IAV-HA (H3SKE, 1:100), mouse anti IAV-HA (A1, 1:1500), rabbit anti IAV-HA (Pinda, 1:5000). Rabbit anti-Lamp1 (9091s, 1:500), rabbit anti-His tag (12698s, 1:400), rabbit anti-EEA1 (3288s, 1:300) from Cell Signalling Technology; mouse anti-EEA1 (610457, 1:300), mouse anti-GM130 (610823, 1:300), mouse anti-CD63 (556019, 1:500) from BD Biosciences; rabbit anti-FAM21 (PA5-35883, 1:350), rabbit anti-WASH1 (PA5-51731, 1:100) from Invitrogen; rabbit anti-GLUT1 (AB115730, 1:500), rabbit anti-EGFR (AB52894, 1:350), rabbit anti-ARPC5 (AB51243, 1:100) from Abcam; mouse anti-VPS35 (374372, 1:250) from Santa Cruz Biotechnology; and rat anti-Myc tag (9E1, 1:400) from ChromoTek. Secondary antibodies used were goat anti-mouse IgG (H+L) Cross-Adsorbed Secondary Antibody, Alexa Fluor™ 488 (A-11001, 1:1000), Alexa Fluor™ 568 (A-11004, 1:500) and Alexa Fluor™ 647 (A-21235, 1:500), goat anti-rabbit IgG (H+L) Cross-Adsorbed Secondary Antibody, Alexa Fluor™ 488 (A-11008, 1:1000), Alexa Fluor™ 568 (A-11011, 1:500) and Alexa Fluor™ 647 (A-21244, 1:500), and goat anti-rat IgG (H+L) Cross-Adsorbed Secondary Antibody, Alexa Fluor™ 568 (A-11077, 1:500).

### IAV entry assays

IAV entry assays were performed as previously described^28,52^. Briefly, for IAV binding assay, cells were incubated with 0.25 µl of 1.95×10^9^ infectious units/ml of IAV X-31 per well of 96-well plate for 45 min at 4 °C. After incubation, cells were washed with cold infection medium twice and fixed with 4% formaldehyde, and IIF was performed with anti-HA antibody (Pinda), hoechst, and phalloidin labelled with Alexa Fluor^TM^ 568 (Invitrogen, A12380).

For IAV endocytosis assay, control cells were pretreated with 25 µg/ml chlorpromazine (Cpz, Merck, C8138) for 10 min at 37 °C. After pretreatment, cells were incubated with 0.25 µl of 1.95×10^9^ infectious units/ml of IAV X-31 per well of 96-well plate for 30 min at 37 °C. In control wells, Cpz was also added during virus incubation. After 30 min of incubation, cells were washed with PBS and fixed with 4% formaldehyde. After fixation, cells were washed with PBS and blocking solution (BS, 5% FBS and 1% BSA in 1x PBS) was added for 1 h. Cells were incubated with rabbit HA (Pinda, 1:1000) in BS at 4 °C overnight to block non-internalized HA epitopes. After incubation, cells were washed three times with PBS for 10 min each, followed by incubation with a secondary anti-rabbit AF647 conjugate. The cells were washed three times with PBS and fixed again with 4% formaldehyde to fix the secondary antibody bound to Pinda. Cells were then permeabilized with PS for 30 min and incubated with mouse HA1 (1:100) for 2 h at RT which will recognize endocytosed HA epitopes. After incubation, cells were washed with PBS and incubated with secondary anti-mouse AF488 conjugate and hoechst and phalloidin labelled with Alexa Fluor^TM^ 568. Cells were washed again with PBS and imaging was performed at 40x in CQ1.

For HA acidification and M1 uncoating assays, cells were incubated with 0.25 µl of 1.95×10^9^ infectious units/ml of IAV X-31 per well of 96-well plate for 60 min and 120 min at 37 °C, respectively. In control wells, BafA1 (50 nM) was also added during virus incubation. After incubation, cells were washed with PBS and fixed with 4% formaldehyde. For vRNP nuclear import assay, cells were incubated with 0.25 µl virus of 1.95×10^9^ infectious units/ml of IAV X-31 per well of 96-well plate for 4 h as described above. Cells were washed with PBS and fixed with 4% formaldehyde. IIF was performed to detect the acid conformation of HA using an acid conformation-specific antibody (A1) for HA acidification assay, and an anti-M1 antibody (HB-64) for M1 uncoating assay.

For IAV fusion assay, cells were incubated with 1 µl labelled virus per well for 45 min at 4 °C. After virus binding, cells were washed with cold infection medium and shifted to 37 °C for 90 min. Cells were then washed with PBS and fixed with 4% formaldehyde. For labelling X-31, lipophilic dyes SP-DiOC18 (Invitrogen, D7778) and R18 (Invitrogen, O246) were dissolved in absolute ethanol at a working concentration of 33 µM and 67 µM, respectively. After the preparation of the dye mixture, 10 µl of IAV X-31 (1.95×10^9^ infectious units/ml) is diluted in 140 µl of PBS and then 1.8 µl of this dye mixture is added to the diluted virus with constant low speed vortexing. The virus is allowed to get labelled for 60 min at 4 °C with continuous rotation at low speed. After the virus is labelled, it is diluted with infection medium and filtered through a hydrophobic filter before use.

### IAV plasma membrane-bypass uncoating assay

IAV plasma membrane-bypass uncoating was performed as previously described^29^. Briefly, cells were seeded in a 96-well plate. After the cells reached ∼90% confluency, plasma membrane-bypass assay was performed. Cells were incubated with 1 µl of 1.95×10^9^ infectious units/ml of IAV X-31 per well of 96-well plate for 45 min at 4 °C. After binding, the virus-containing medium was discarded and the cells were washed with cold infection medium. Stop medium (50 mM HEPES pH7.4, 20 mM NH_4_Cl in DMEM) was added in control wells. In WT and KO/KD cells, fusion media (50 mM citrate buffer in DMEM, pH 5.0) was added and the plate was shifted to 37 °C in CO_2_ incubator for 2 min. Immediately after 2 min, the fusion medium was removed very gently and stop media was added. The plate was placed in CO_2_ incubator at 37 °C for 5 min. Cells were fixed with 4% formaldehyde. IIF was performed against IAV M1 (HB64, 1:30) and imaging was performed at 40x in CQ1.

### EGF, transferrin, and cholera toxin B uptake assays

For EGF uptake, transfected cells were serum-starved for 20 h in plain DMEM. Control wells were pre-treated with Cpz for 10 min. Alexa Fluor™ 488-EGF (30 ng/ml) (Invitrogen, E13345) was added, and cells were incubated for 30 min at 37 °C in a CO₂ incubator. Cpz was maintained during EGF incubation in control wells. After incubation, plates were shifted to 4 °C, washed with ice-cold PBS, treated with ice-cold acid wash buffer (150 mM NaCl, 50 mM Glycine, pH-3.0) for 2 min, washed again with PBS, and fixed with 4% formaldehyde at room temperature. Cells were counterstained with Hoechst and imaging was performed.

For transferrin (Tfn) uptake, transfected cells were serum-starved for 4 h in plain DMEM. siCLTC was used as a positive control. Alexa Fluor™ 488-Tfn (30 µg/ml in serum-free media) (Invitrogen, T13342) was added and allowed to bind at 4 °C for 45 min. Cells were then washed three times with ice-cold PBS, shifted to 37 °C in serum-free media for 10 min, and immediately returned to 4 °C to stop uptake. After washing with ice-cold PBS, cells were treated with acid wash buffer for 2 min, washed again, and fixed with 4% formaldehyde at RT, followed by Hoechst staining and imaging.

For cholera toxin B (CTxB) uptake, transfected cells were serum-starved for 2 h in plain DMEM. Control wells were pre-treated with MβCD (10mM) (Merck, C4555) for 1 h. Alexa Fluor™ 488-CTxB (750 ng/ml in serum-free media) (Invitrogen, C34775) was added and allowed to bind at 4 °C for 45 min. After binding, cells were washed three times with ice-cold PBS, then incubated in serum-free media at 37 °C for 10 min or 30 min. MβCD was maintained during binding and uptake in control wells. Plates were then shifted to 4 °C, washed with ice-cold PBS, treated with acid-wash buffer for 2 min, washed again with PBS, and fixed in 4% formaldehyde at RT. IIF was carried out to stain the Golgi complex with anti-GM130.

### Dextran uptake assay

For dextran (Dex) uptake, control wells were pre-treated with 50 µM EIPA for 30 min at 37 °C. Plate was incubated at ice for 10 min to stop endocytosis pathways. FITC-labelled Dex (Invitrogen, D1822) was diluted to 100 µg/ml in serum-free media, sonicated and added to the cells, followed by incubation for 30 min at 37 °C in a CO₂ incubator. In control wells, EIPA was maintained during uptake. After incubation, the plate was shifted to ice to stop uptake, and cells were washed thrice with PBS. Cells were then fixed with 4% formaldehyde at RT, counterstained with Hoechst, and imaging was performed.

### DENV-2 infection assay

HepG2 cells were reverse transfected with 10 nM siRNAs using Lipofectamine RNAiMAX and incubated for 72 h at 37 °C in CO_2_ incubator. After incubation, cells were infected with DENV-2 for 48 h. The virus was diluted in infection medium (2% BSA in DMEM). BafA1 was used as an inhibitor of virus infection. Following 2 h of endocytosis, the virus-containing medium was removed and replaced with maintenance medium (DMEM supplemented with 2% FBS). After 48 h, cells were washed with PBS and fixed with 4% formaldehyde for 20 min at RT and then IIF was performed to detect viral envelope protein (E). Cells were imaged using CQ1. E protein intensity was quantified using ImageJ software and average intensity per cell was calculated.

### Western blotting

For protein detection, cells were lysed using cell lysis buffer (150 mM NaCl, 50 mM Tris pH 7.4, 1% Nonidet P-40, 0.1% SDS and 0.5% deoxycholate, protease inhibitor cocktail). The lysate was incubated on ice for 30 min with vortexing every 10 min followed by centrifugation at 17000 g for 20 min. The supernatants were collected and mixed with 6x SDS loading dye. Samples were boiled at 95 °C for 10 min and separated on 12.5% SDS-PAGE at 90V for 2 h. Separated proteins were transferred to nitrocellulose membrane at 90V for 1.5 h followed by membrane blocking with 5% skimmed milk or 5% BSA in 0.1% Tween in 1x Tris-buffered saline (TBST) for 1.5 h. Membrane was incubated with primary antibody overnight at 4 °C. The membrane was washed with 0.1% TBST for 5 min, three times at RT, and incubated with an HRP-tagged secondary antibody for 1 h at RT followed by three washes for 10 min each. The signal was detected using Clarity™ Western ECL Substrate (Bio-Rad, 170-5060).

Primary antibodies used were mouse anti IAV-NP (HB65, 1:50), mouse anti IAV-M1 (HB64, 1:30) from ATCC; rabbit anti-GAPDH (2118s, 1:2000), mouse anti-Lamp1 (15665s, 1:1000), mouse anti-PCNA (2586s, 1:1000) from Cell Signalling Technology; rabbit anti-FAM21 (PA5-35883, 1:350), rabbit anti-WASH1 (PA5-51731, 1:100) from Invitrogen; mouse anti-VPS35 (374372, 1:250), mouse anti-His tag (sc-53073, 1:1000) from Santa Cruz Biotechnology; rat anti-Myc tag (9E1, 1:400) from ChromoTek, mouse anti-Myc-tag (M047-3) from MBL; rabbit anti-GFP purified antibody from Abgenex (1:2500); rabbit anti-CCDC53 (24445-1-AP, 1:500) from Proteintech; rabbit anti-SWIP (A304-919A-T, 1:2000) from Bethyl Laboratories; mouse anti-FLAG (F1804, 1:1000) from Merck. Secondary antibodies used were anti-mouse IgG, HRP-linked antibody (7076s, 1:2000), anti-rabbit IgG, HRP-linked antibody (7074s, 1:2000), and anti-rat IgG, HRP-linked antibody (7077s, 1:2000) from Cell Signalling Technology.

### Co-immunoprecipitation assay

HEK293T cells were transfected with the plasmids expressing proteins of interest. After 12-14 h, cells were lysed with ice cold Lysis buffer (10 mM Tris/Cl pH 7.5, 150 mM NaCl, 0.5 mM EDTA, 0.5% NP-40). Lysate was incubated on ice for 30 min, with extensive pipetting at every 10 min. Lysate was then centrifuged at 17,000 g for 10 min at 4 °C. The supernatant was collected and 5% of it was taken as input fraction. The remaining supernatant was incubated with Myc-Trap® (Proteintech, yta) or GFP-Trap® Agarose beads (Proteintech, gta) or anti-DYKDDDDK Tag (L5) Affinity Gel (FLAG-Trap, BioLegend, 651501) for 2 h at 4 °C with rotation at 13 rpm. After incubation, antigen-conjugated agarose beads were washed three times with ice-cold wash buffer (10 mM Tris/Cl pH 7.5, 150 mM NaCl, 0.05% NP40, 0.5 mM EDTA). The protein complexes were then eluted from the beads by boiling them at 95 °C for 10 min. Beads were again centrifuged at 2000 rpm for 1 min and the supernatant was loaded onto 12.5% SDS-PAGE gel. Western blotting was performed with specified antibodies.

### Confocal and high-resolution imaging

For confocal imaging, cells were transfected or seeded on coverslips (VWR®, 631-0148) and experiments were performed. Cells were then fixed with 4% formaldehyde for 20 min at RT and IIF was performed as previously described and then the coverslips were mounted on glass slides using Fluoromount-G® (Southern Biotech, 0100-01). The slides were allowed to dry overnight and then the imaging was performed under a 63x oil immersion lens in a Leica SP8 upright confocal microscope. Z-stack images were acquired with 0.3 µm interval and later stacked to analyze the data using ImageJ software. For GLUT1 localization, HeLa cells were reverse transfected with 20 nM siTRIM62. Appropriate controls were included. After 72 h of transfection, the control wells were treated with brefeldin A (1 µM) (Merck, B6542) or monensin (1x) (Invitrogen, 00-4505-51) for 1 h and cells were fixed with 4% formaldehyde for 20 min at RT.

High-resolution imaging was done using 63x oil immersion lens on ZEISS LSM 980 Elyra 7 super-resolution microscope equipped with a monochrome cooled high-resolution AxioCamMRm Rev. 3 FireWire(D) camera with Airyscan 2.0 processing. To detect co-localization of TRIM62-Myc with WASH1 or FAM21, A549 cells were seeded on coverslips and transfected with plasmid encoding TRIM62-Myc. After 16 h, cells were treated with 1 µM apilimod (Merck, SML2974) and incubated at 37 °C for 40 min in CO_2_ incubator. Cells were then fixed with 4% formaldehyde. IIF was performed to detect TRIM62-Myc and endogenous WASH1 or FAM21 and imaging was performed.

### IAV plaque assay

A549 WT and TRIM62 KO cells were seeded in a 6 well plate. At 60% confluency, cells were infected with IAV WSN (H1N1) at an MOI of 0.01 for 24 h at 37 °C in CO_2_ incubator. After 24 h, the cell supernatant was collected and centrifuged at high speed to remove cell debris. This cleared supernatant was collected and diluted to 10^1^, 10^2^, 10^4^, and 10^6^ times in infection medium and then added to the monolayer of MDCK cells seeded in a 12 well plate for 2 h and cells were incubated at 37 °C in CO_2_ incubator. After 2 h, cells were washed with infection medium and then overlaid with warm infection medium containing 0.6% low melting agarose (Genaxy, GEN-LE-100), 1x penicillin-streptomycin and 1 µg/ml TPCK-Trypsin (Merck, T1426). Agarose was allowed to solidify for 15 min at RT and then plates were shifted to 37 °C in CO_2_ incubator for 48 h. Plaques were observed and cells were fixed with 8% formaldehyde for 60 min at RT. After fixation, the agarose plugs were carefully removed without disturbing the cells and were stained with crystal violet blue for 15 min with constant shaking at low speed. Plates were then washed with water to remove extra stain. Plaques were counted and plaque forming units (PFU)/ml was calculated.

### Live cell imaging

For live-cell imaging, A549 cells were grown in 35 mm cell culture dishes (ibidi, 81156) and transfected with expression plasmids using Lipofectamine™ LTX Reagent with PLUS™ Reagent. After 24 h, videos were recorded using 63x oil immersion lens in Zeiss LSM 980 Airyscan 2.0 equipped with CO_2_ gas chamber maintained at 37 °C. For apilimod treatment, cells were treated with 1 µM apilimod and incubated at 37 °C in CO_2_ incubator for 40 min and then videos were recorded.

### Proximity ligation assay (PLA)

Cells were transfected with siControl and siTRIM62 in a 96-well plate and fixed with 4% formaldehyde after 72 h of transfection. PLA was performed on fixed cells using Duolink® In Situ Red Starter Kit (Merck, DUO92101) according to the manufacturer’s protocol. All incubations were done at 37 °C in humidified chamber. In summary, blocking solution was added to the wells and the plate was incubated for 1 h. After blocking, primary antibodies were diluted in antibody-diluent, added in the wells, and incubated for 2 h. After incubation, cells were washed with wash buffer A for 10 min each and anti-rabbit plus and anti-mouse minus probes were added for 1 h. Cells were washed with wash buffer A and incubated with ligase for 30 min, followed by washing. In the last step, polymerase was added and the plate was incubated for 100 min. Cells were washed with wash buffer B for 10 min twice and the last washing was done with 0.01x wash buffer B for 1 min. Hoechst was diluted in PBS and cells were stained and imaged in CQ1.

For TRIM62-Myc and ΔP-S TRIM62-Myc, A549 cells stably expressing the plasmids were seeded in 96-well plate and fixed with 4% formaldehyde and PLA was performed.

### Yeast 2-hybrid (Y2H) assay

To check direct protein-protein interaction, TRIM62 was cloned in the pGAD-T7 vector and the suspected partner is cloned in the pGBK-T7 vector. *Saccharomyces cerevisiae* Y2HGold cells were made competent using a modified LiOAc method. 200 ng DNA of each of the two plasmids were transformed together in yeast competent cells by adding six-fold volume of 40% filtered polyethylene glycol (PEG, Merck, P4338). This suspension was mixed well and incubated at 30 °C for 30 min. Following this incubation, cells were shifted at 42 °C for 25 min. Cells were then plated onto plates containing synthetic medium lacking the amino acid leucine and tryptophan and incubated at 30 °C for 72 h until growth was observed. Single colonies of transformed yeast cells are selected, diluted in 100 µl of filtered autoclaved water. Then, a six-fold serial dilution of transformed colonies was spotted in both control plate (- leucine -tryptophan) and the selection plate (-leucine -tryptophan -histidine). Plates were incubated at 30 °C for 72-96 h till growth was observed. Growth on plate containing -histidine selection media indicated interaction between two proteins.

### Image processing and analysis

All images at 20x and 40x objectives were acquired using CQ1. Experiments were carried out in 96-well optical-bottom Greiner plates. The focal plane was set automatically using Hoechst as a reference. Z-stacks were collected within a 20 µm range of the fixed center. Nine random fields were selected from one well, and the same fields were imaged across all sample wells. Maximum intensity projection (MIP) images were generated in TIFF format for each laser channel. For quantifying infection percentage, image analysis pipelines were built in CellProfiler (v2.2.0) and KNIME (v3.7.2) as previously described^53^.

For NP intensity, ImageJ (2.0.0-rc-69/1.52p) was used. Nuclei were thresholded and counted, and NP signals were thresholded for particle analysis. Thresholded area × mean intensity was normalized to the cell number to calculate NP intensity per cell. Data were plotted in GraphPad Prism 9, and statistical analysis was done using one-way ANOVA with multiple comparisons. IAV entry assay intensity data were processed similarly.

For IAV binding assays, surface-bound HA protein, F-actin, and nuclei were stained. F-actin marked the cellular area, which was Gaussian-blurred and thresholded. This mask was applied on the HA channel to extract HA signal. HA intensity for endocytosis and acidification assays was calculated in the same way. For M1 uncoating, cells with dispersed M1 were counted as positive, and percentage uncoating was calculated relative to total nuclei. For dispersed M1 intensity, a 40 × 40 square ROI was drawn in the cytoplasm, and mean intensity was measured. For vRNP nuclear import, cells positive for nuclear NP were scored, and percentage was calculated. To quantify nuclear NP intensity, a nuclear mask was created from Hoechst and applied to the NP channel.

For co-localization analysis, images were acquired at 63x oil immersion on the Leica SP8 upright confocal microscope at 1024 × 1024 resolution with z-stacks. Mander’s coefficients were calculated using the JaCoP plugin in ImageJ. TRIM62-Myc and WASH1 or FAM21 co-localization upon apilimod treatment was imaged using ZEISS LSM 980 Elyra 7 in super-resolution mode. Images were converted to 8-bit before analysis. Intensity-distance profiles were generated by drawing a line across co-localized regions and plotting signal intensity changes using the profile tool in Zen 3.4 Blue software.

### Statistical analysis

All data are presented as mean ± standard deviation (SD) from at least three independent experiments (n ≥ 3). Statistical analyses were performed using GraphPad Prism 9. Differences between groups were assessed using either one-way ANOVA with multiple comparisons or an unpaired t-test, as appropriate. A p-value of < 0.05 was considered statistically significant.

## Data and code availability

All raw data to reanalyse the data reported in this paper are available on request. This manuscript does not report original code. The lead contact can provide any additional information related to the data reported in this paper.

## Acknowledgements

We thank many past and present members of the Banerjee laboratory for useful discussions and intellectual inputs. We thank IISER Mohali, Department of Biotechnology (DBT), Govt. of India (grant no. BT/PR38441/MED/29/1498/2020), Department of Science and Technology, Govt. of India (FIST grant no. SR/FST/LS-II/2017/97 to the Department of Biological Sciences, IISER Mohali) for financial support. We also thank the Council of Scientific and Industrial Research (CSIR), University Grants Commission (UGC), DBT, Govt. of India, for providing fellowships to K.G., S.P., G.K. The authors acknowledge the support of Dr. Mahak Sharma (IISER Mohali) for providing the Y2HGold strain of *Saccharomyces cerevisiae* and Dr. Shravan Kumar Mishra (IISER Mohali) and his lab members for helping us with Y2H experiments. We extend our sincere thanks to Dr. Amit Tuli (CSIR-IMTECH), Dr. Pradeep Uchil (Yale University, USA), Dr. Debojyoti Chakraborty (IGIB, India), Dr. Santosh Chauhan (CCMB Hyderabad, India), Prof. Juan S. Bonifacino (NIH, USA), Prof. Lenka Libusova (Charles University, Czech Republic), Prof. Alexis Gautreau (CNRS, France), and Prof. Peter J. Cullen (University of Bristol, UK) for providing us with different plasmids used in this study.

## Author contributions

Conceptualization, funding acquisition and supervision, I.B.; intellectual inputs throughout the study, I.B. and K.G.; experiments and data acquisition, K.G., S.P., R.B., T.P. and G.K.; formal analysis, K.G.; validation, K.G.; writing – original draft, K.G.; writing – review and editing, I.B. and K.G.; all authors reviewed and accepted the final contents of the manuscript.

## Supplementary information

Supplementary Fig. 1-8 and Supplementary Tables 1-3

