## Supplementary Information for "TRIM62 promotes lipid raft-mediated entry of influenza A virus by regulating WASH localization"

**Supplementary Fig. 1**

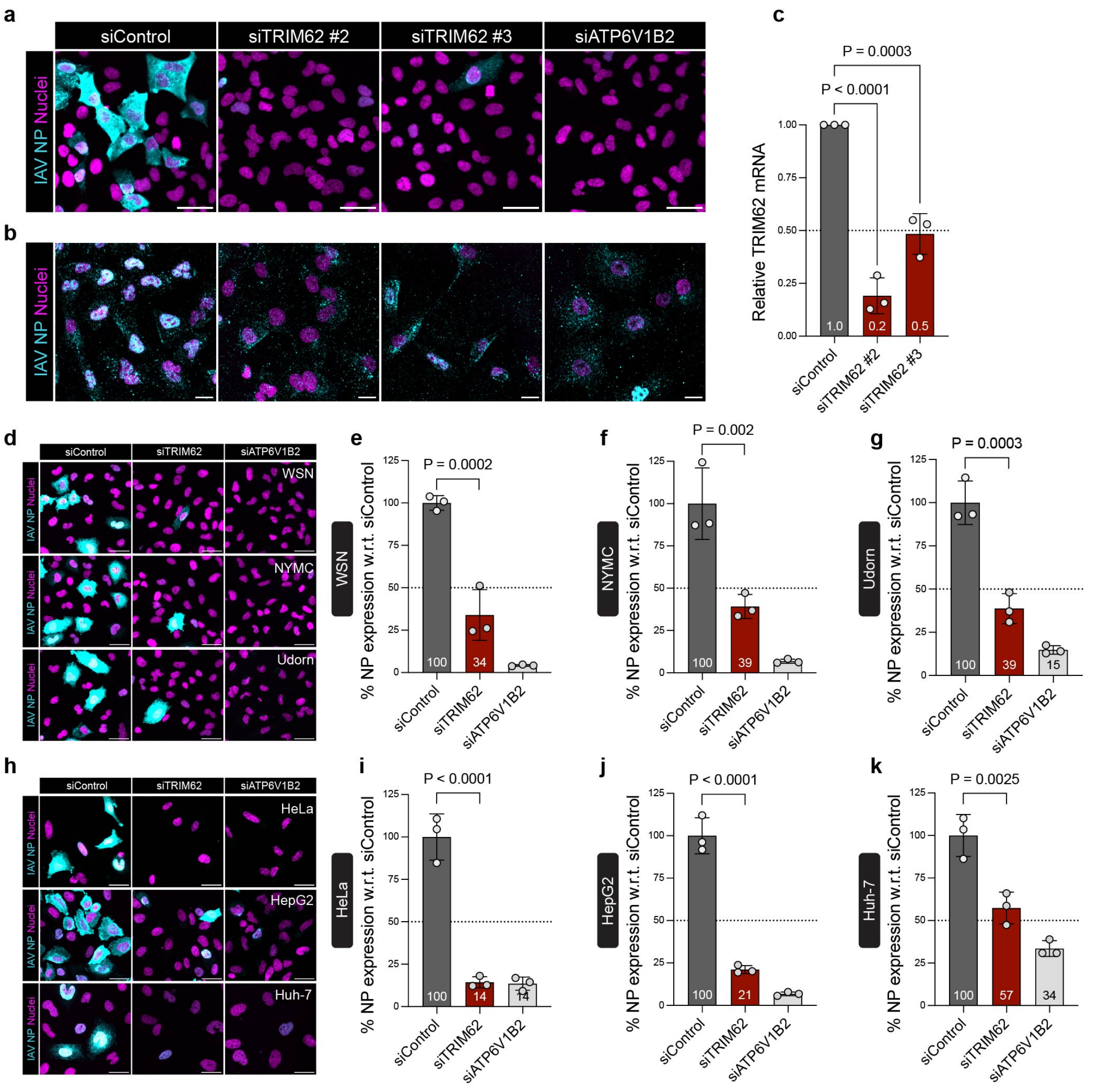


**Supplementary Fig. 1 Knockdown of TRIM62 effectively reduces *TRIM62* mRNA levels and IAV infection across different viral strains and cell lines.**

**a,b** High-content confocal images, showing IAV X-31 infection (**a**) and vRNP nuclear import (**b**) at 10 h.p.i. and 4 h.p.i., respectively. NP (cyan) and nuclei (magenta). Scale bar, 20 µm. **c** Quantification of mRNA levels of *TRIM62* upon knockdown using two independent siRNAs targeting TRIM62. **d-g** High-content confocal images (**d**) and quantification of IAV infection in control and TRIM62 KD cells infected with WSN (**e**), NYMC (**f**), and Udorn (**g**) for 10 h. NP (cyan), and nuclei (magenta). Scale bars, 20 µm. $n$ > 4000 cells per sample. **h-k** High-content confocal images (**h**) and quantification of IAV infection in HeLa (**i**), HepG2 (**j**), and Huh-7 (**k**) cells infected with X-31 strain for 10 h. Scale bars, 20 µm. $n$ > 1500 cells per sample. Data in **c**,**e-g**,**i-k** were analysed using one-way ANOVA using multiple-comparisons, P values are indicated. P values < 0.05 were considered significant. All bar graphs show the mean of n = 3 ± SD.

#### Supplementary Fig. 2

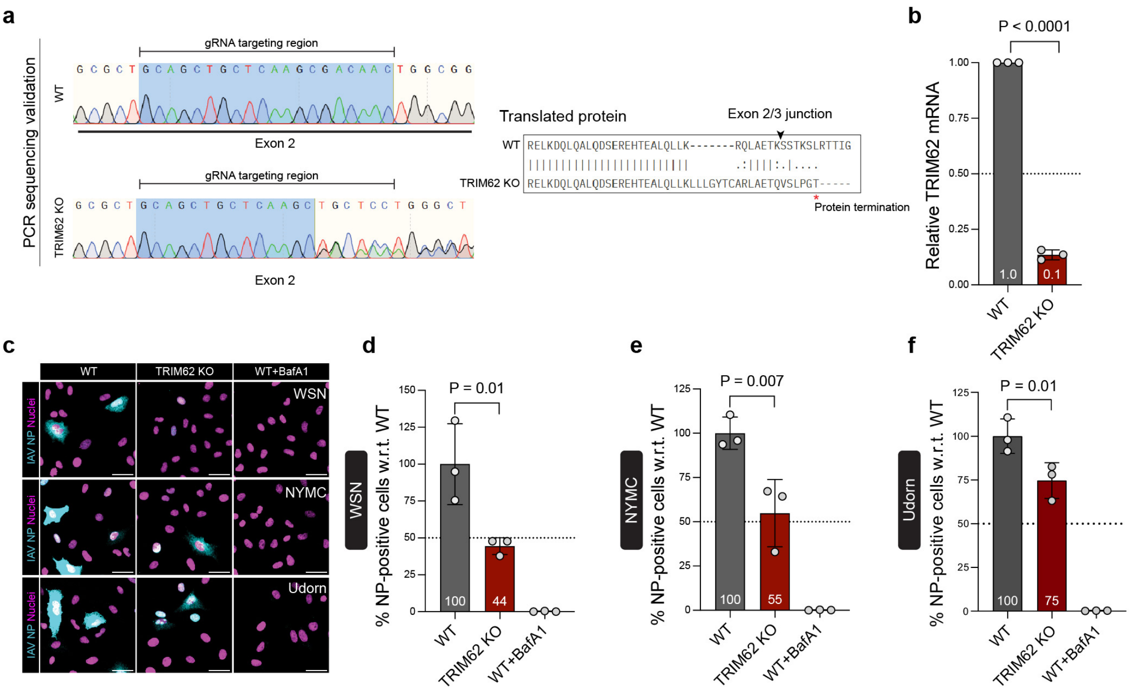


**Supplementary Fig. 2 TRIM62 KO validation by sanger sequencing and RT-PCR, and reduction in IAV infection in TRIM62 KO cells across different viral strains.**

**a** Chromatogram (left) showing DNA sequence downstream of the gRNA-targeted genomic region from WT and TRIM62 KO cells. gRNA targeting region in exon 2 is highlighted. DNA sequences from WT and TRIM62 KO are translated using Expasy Translate tool and pairwise aligned using EMBOSS Needle tool (right), indicated protein termination in KO. **b** Quantification of mRNA levels of *TRIM62* in TRIM62 KO cells compared to WT. **c-f** High-content confocal images (**c**) and quantification of IAV infection in TRIM62 KO and WT cells infected with WSN (**d**), NYMC (**e**), and Udorn (**f**) for 10 h. Scale bars, 20 µm. *n* > 3500 cells per sample. Data in **b** was analyzed using unpaired two-tailed *t*-test. Data in **d-f** were analysed using one-way ANOVA using multiple-comparisons, P values are indicated. P values < 0.05 were considered significant. Bar graph shows the mean of n = 3 ± SD, and all data are representative of three biological replicates (N = 3).

### Supplementary Fig. 3

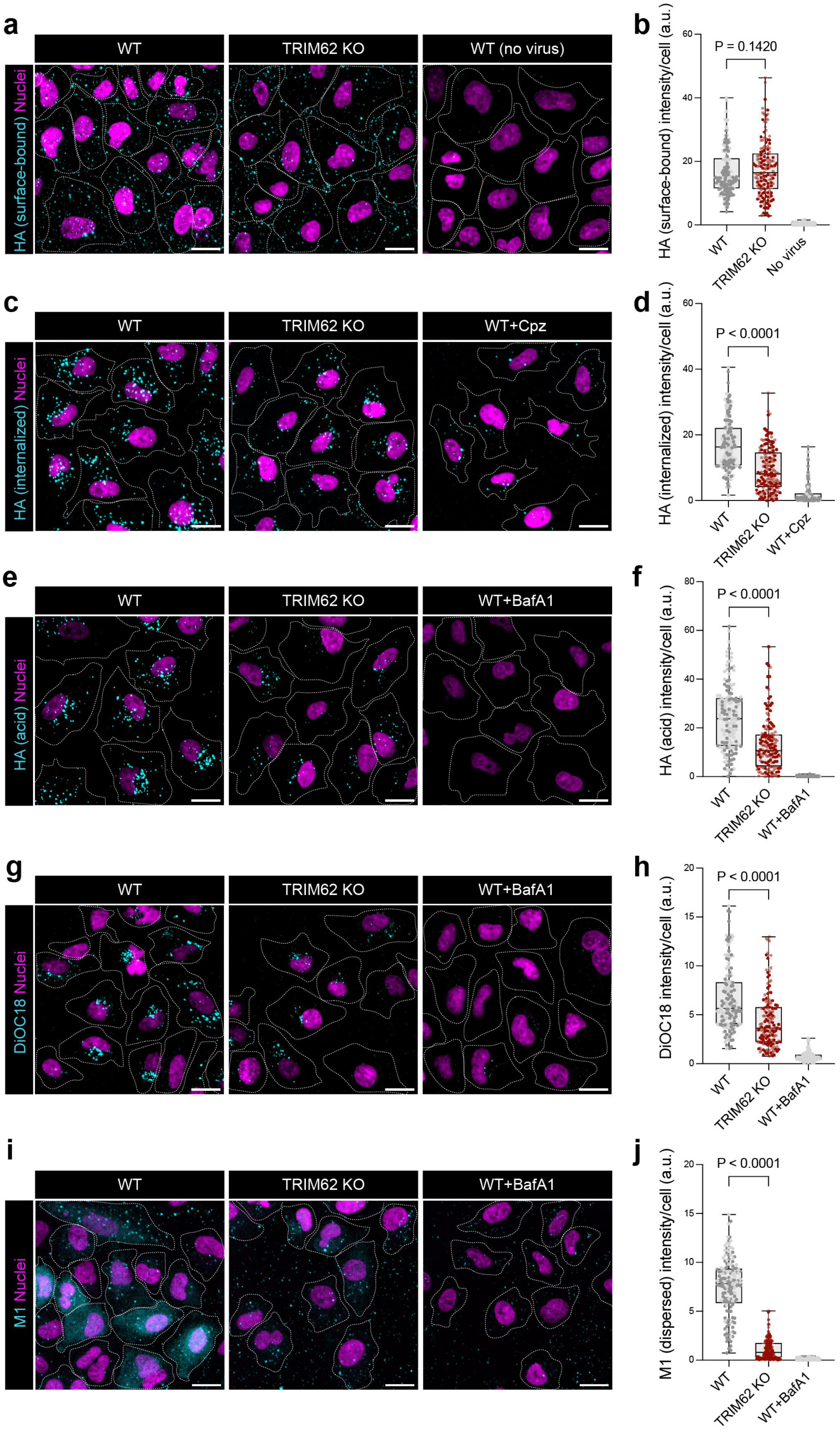

**Supplementary Fig. 3 TRIM62 deficiency blocks IAV endocytosis and downstream entry steps.**

**a,b** High-content confocal images (**a**) and intensity analysis (**b**) of surface-bound IAV particles detected by IIF against HA. HA (cyan) and nuclei (magenta). Cell boundaries are outlined. Scale bars, 20 µm. *n* = 158 cells per sample. **c,d** High-content confocal images (**c**) and intensity analysis (**d**) of internalized IAV particles (HA) at 30 min post-infection. HA (cyan) and nuclei (magenta). Scale bars, 20 µm. *n* = 156 cells per sample. **e,f** High-content confocal images (**e**) and intensity analysis (**f**) of acidified HA at 1 h.p.i. Scale bars, 20 µm. *n* = 146 cells per sample. **g,h** High-content confocal images (**g**) and intensity analysis (**h**) of DiOC18 labelled X-31 upon IAV fusion at 1.5 h.p.i. DiOC18 (cyan) and nuclei (magenta). Scale bars, 20 µm. *n* = 152 cells per sample. **i,j** High-content confocal images (**i**) and intensity analysis (**j**) of cytosolic M1 (dispersed) upon M1 uncoating at 2 h.p.i. M1 (cyan) and nuclei (magenta). Scale bars, 20 µm. *n* = 150 cells per sample. Data in **b**,**d**,**f**,**h**,**j** were analysed using one-way ANOVA using multiple-comparisons, P values are indicated. P values < 0.05 were considered significant. All data are representative of three biological replicates (N = 3).

#### Supplementary Fig. 4

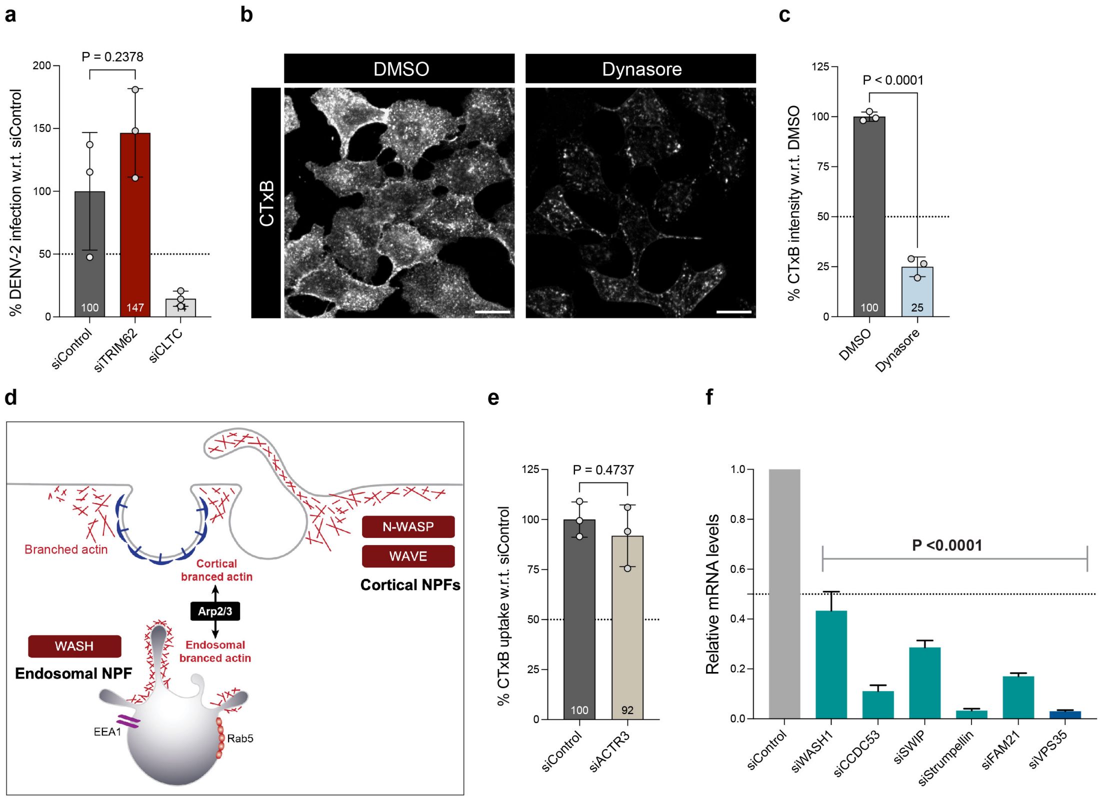


**Supplementary Fig. 4 CTxB uptake is dynamin-dependent, but branched actin independent in A549 cells.**

**a** Quantification of DENV-2 infection (% E expression at 48 h.p.i., normalized) in HepG2 cells. n >1000 cells per sample. **b,c** High-content confocal images (**b**) and quantification (**c**) of CTxB uptake in DMSO- and dynasore-treated cells at 10 min. *n* >1700 cells per sample. **d** Schematic representation of branched F-actin nucleation by Arp2/3 complex and cortical or endosomal NPFs. **e** Quantification of CTxB uptake in ACTR3-depleted cells at 30 min. *n* >2000 cells per sample. **f** Validation of siRNA KD efficiency against WASH subunits and VPS35 used in the study by RT-PCR. Quantification of mRNA levels of target genes in siRNA-treated cells. Data in **c,e** were analyzed using unpaired two-tailed *t*-test. Data in **a,f** were analysed using one-way ANOVA using multiple-comparisons, P values are indicated. P values < 0.05 were considered significant. Bar graph shows the mean of n = 3 ± SD, and all data are representative of three biological replicates (N = 3).

#### Supplementary Fig. 5

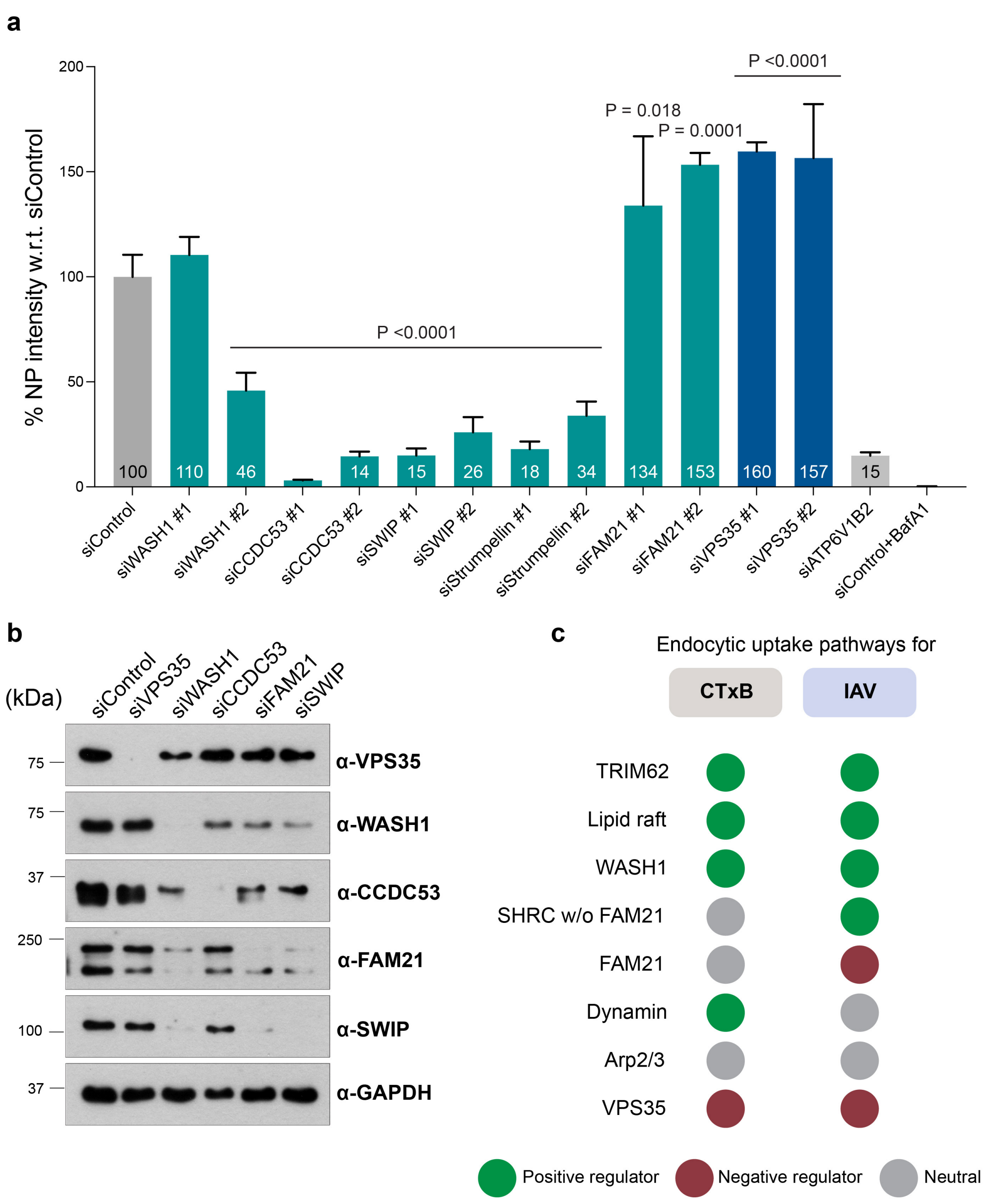


**Supplementary Fig. 5 Functional antagonism between the WASH complex and VPS35, and differential requirements in IAV and CTxB uptake.**

**a** Quantification of IAV infection (% NP expression at 10 h.p.i., normalized) in A549 cells infected with IAV X-31 strain (MOI = 0.01) in an RNAi screen targeting VPS35 and WASH complex subunits. Data shown for two independent siRNAs targeting each gene. *n* >2000 cells per sample. **b** Western blots showing the effects of KD of individual WASH subunits and

VPS35 on the protein levels of the others. **c** Schematic showing the similar/differential requirements of TRIM62, WASH complex subunits, VPS35, dynamin, and Arp2/3 in the lipid raft-mediated uptake of CTxB and IAV. Data in **a** was analysed using one-way ANOVA using multiple-comparisons, P values are indicated. P values $< 0.05$ were considered significant.

#### Supplementary Fig. 6

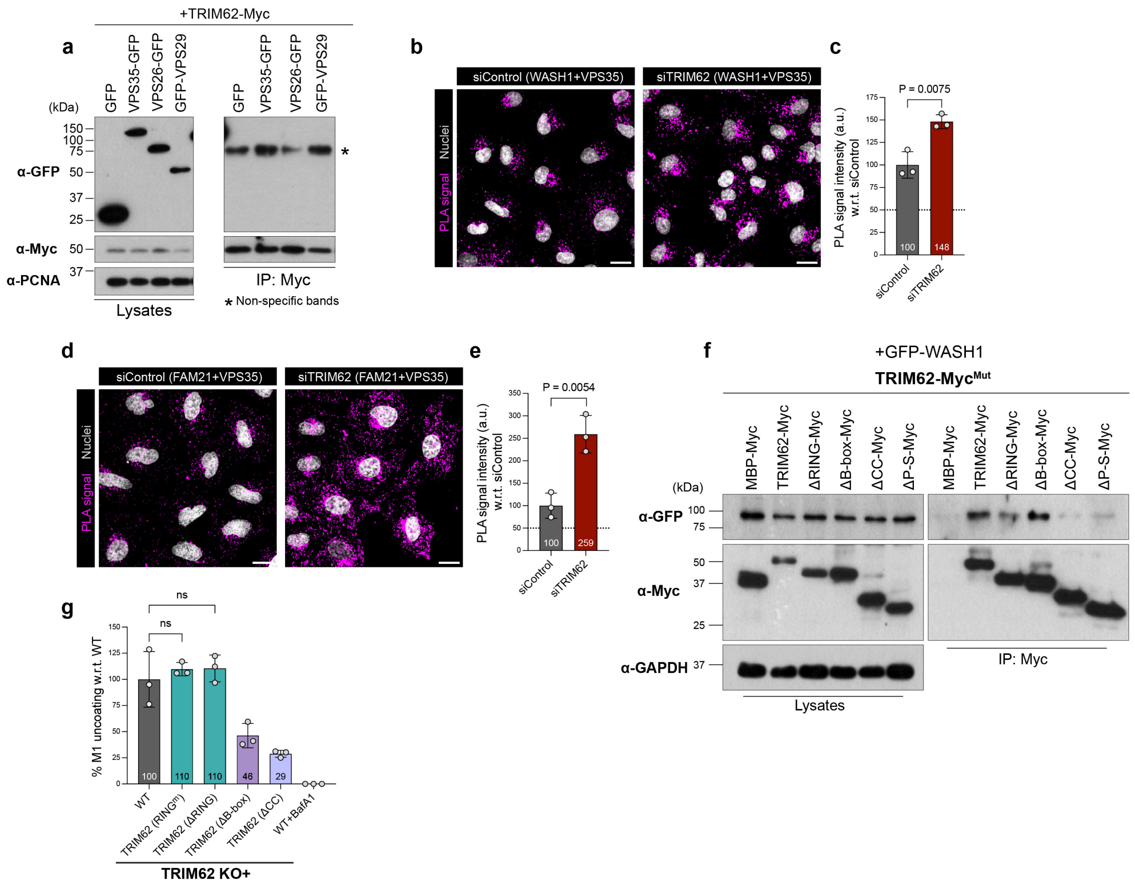


**Supplementary Fig. 6 TRIM62 interacts with the WASH complex, regulates WASH-VPS35 association, and promotes IAV entry independent of its E3 ubiquitin ligase activity.**

**a** HEK293T cells were transfected with TRIM62-Myc and GFP-tagged retromer subunits and lysates were incubated with Myc-Trap beads. Western blotting was done to detect GFP-tagged retromer subunits in the precipitates. PCNA was used as the loading control. * Indicates non-specific bands. **b,c** High-content confocal images (**b**) and quantification (**c**) of PLA signals from VPS35-WASH1 interaction. Scale bars, 20 µm. *n* > 750 cells per sample. **d,e** High-content confocal images (**d**) and quantification (**e**) of PLA signals from VPS35-FAM21 interaction in HeLa cells. Scale bars, 20 µm. *n* > 500 cells per sample. **f** Pull-down assay showing interactions between WASH1 and TRIM62 truncation mutants. HEK293T cells were co-transfected with GFP-WASH1 and TRIM62-Myc mutant constructs, and lysates were incubated with Myc-Trap beads. Western blotting was done to detect GFP-WASH1 in the precipitates. **g** Quantification of % M1 uncoating normalized to WT. *n* > 450 cells per sample. Data in **g** was analysed using one-way ANOVA using multiple-comparisons, P values are indicated. Data in **c,e** were analysed using unpaired two-tailed *t*-test, P values are indicated. P values < 0.05 were considered significant. All bar graphs show the mean of n = 3 ± SD.

**Supplementary Fig. 7**

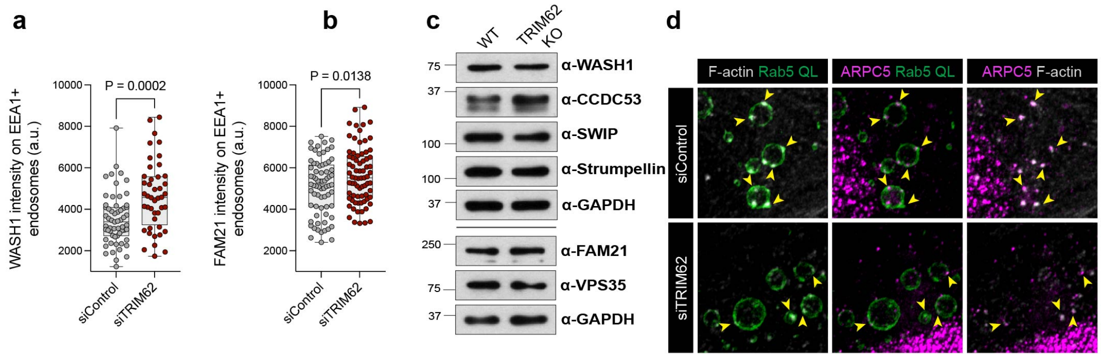


**Supplementary Fig. 7 TRIM62 deficiency enhances WASH endosomal recruitment without altering WASH subunits/VPS35 levels, and impairs WASH1 NPF activity.**

**a,b** Quantification of WASH1 (**a**) and FAM21 (**b**) intensity on EEA1+ endosomes. $n > 45$ cells per sample. **c** Western blots showing the protein levels of WASH subunits and VPS35 in WT and TRIM62 KO cells. **d** Representative confocal microscopy images showing ARPC5 on Rab5QL+ endosomes in control and TRIM62-depleted cells. Data in **a,b** were analyzed using unpaired two-tailed *t*-test, P values are indicated. P values $< 0.05$ were considered significant.

**Supplementary Fig. 8**

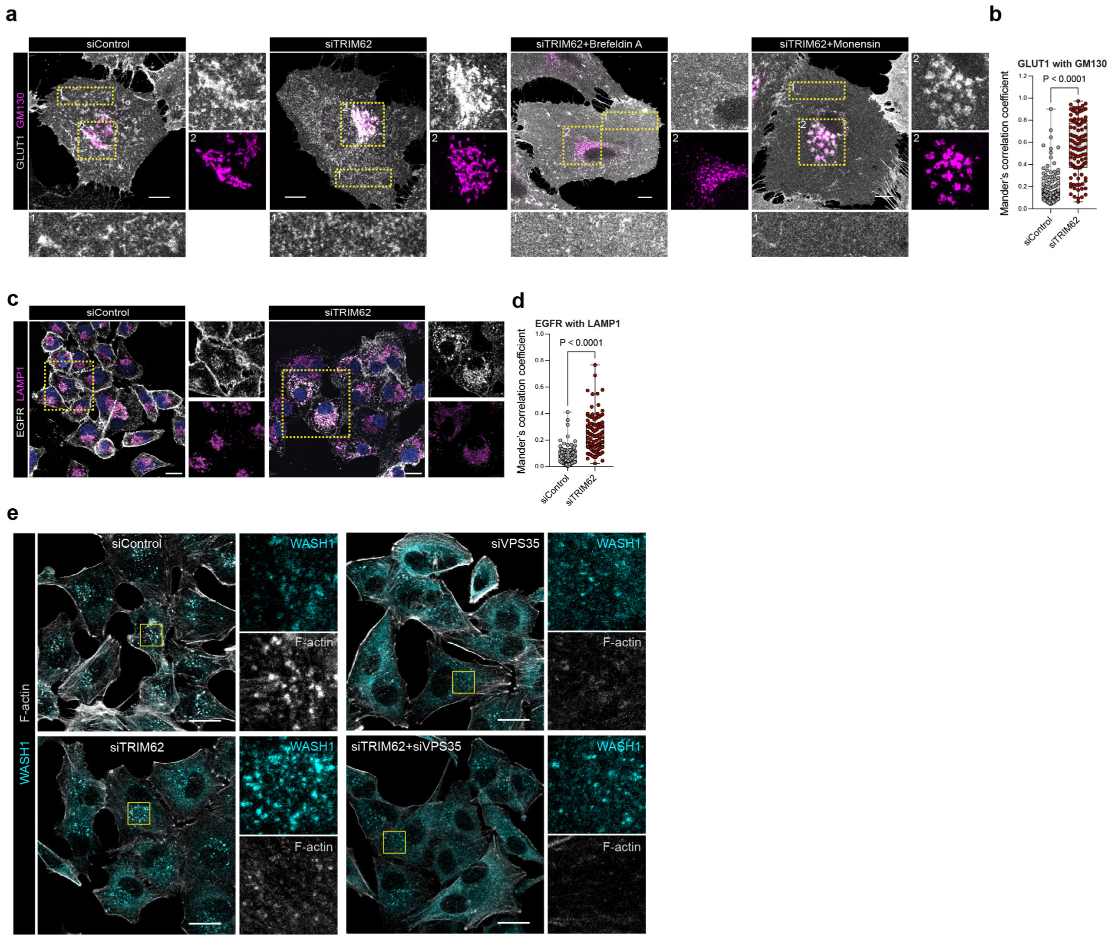


**Supplementary Fig. 8 TRIM62 depletion results in WASH-mediated canonical cargo trafficking defects.**

**a** Representative confocal microscopy images showing GLUT1 (grey) and GM130 (magenta) in HeLa cells with or without the treatment of brefeldin A (1 µM) or monensin (1x) for 1 h. Insets 1 and 2 show GLUT1's surface distribution and Golgi-localization, respectively. Scale bars, 10 µm. **b** Quantification of GLUT1-GM130 colocalization (without brefeldin A or monensin treatment) based on Mander's correlation coefficient. $n$ = 104 cells per sample. **c** Representative confocal microscopy images showing EGFR (grey) and LAMP1 (magenta) in A549 cells. Insets (indicated by yellow boxes) show late-endo/lysosomal distribution of EGFR. Scale bars, 10 µm. **d** Quantification of EGFR-LAMP1 colocalization based on Mander's correlation coefficient. $n$ = 112 cells per sample. **e** Confocal images showing WASH1 (cyan) and F-actin (grey) in control, TRIM62, VPS35, and TRIM62+VPS35 KD cells. Insets (indicated by yellow box) showing zoomed-in images of F-actin patches on WASH1 spots. Scale bars, 20 µm. Data in **b,d** were analyzed using unpaired two-tailed $t$-test, P values are indicated. P values < 0.05 were considered significant.

**Supplementary Table 1. List of plasmids used in this study**

| Plasmid | Source |
|---|---|
| TRIM62-GFP | TRIM62 was PCR amplified from pTRIM62-YFP and inserted into Xba1/BamH1 in pLenti CMV GFP Puro (Addgene #17448) (pTRIM62-YFP was a kind gift from Dr. Pradeep Uchil, Yale University) |
| TRIM62-His | TRIM62 was PCR amplified from pTRIM62-GFP and inserted into Xba1/Sal1 in pLenti CMV GFP Puro. His-sequence was added in primer. |
| TRIM62-His mutants | PCR amplified from pTRIM62-YFP mutants and inserted into Xba1/Sal1 in pLenti CMV GFP Puro (pTRIM62-YFP mutants were kind gifts from Dr. Pradeep Uchil, Yale University) |
| TRIM62-Myc | TRIM62 was PCR amplified from pTRIM62-GFP and inserted into Xba1/Sal1 in pLenti CMV GFP Puro. Myc-sequence was added in primer. |
| TRIM62-Myc mutants | PCR amplified from pTRIM62-His mutants and inserted into Xba1/Sal1 in pLenti CMV GFP Puro. |
| MBP-His (pHLmMBP-6) | Addgene #72346 |
| MBP-Myc | MBP was PCR amplified from pMBP-His and inserted into Xba1/Sal1 in pLenti CMV GFP Puro. Myc sequence was added in primer. |
| GST-Myc | GST was PCR amplified from pGEX-GST and inserted into Xba1/Sal1 in pLenti CMV GFP Puro. Myc sequence was added in primer. (pGEX-GST was a kind gift from Dr. Mahak Sharma) |
| CCDC53-GFP | PCR amplified from cDNA and inserted into Xho1/HindIII in pEGFP-N1 (pEGFP-N1 was a kind gift from Dr. Mahak Sharma) |
| Strumpellin-GFP | PCR amplified from pGFP-Strumpellin and inserted into Nhe1/Xma1 in pEGFP-N1 (pGFP-Strumpellin was a kind gift from Prof. Lenka Libusova, Charles University) |
| FAM21-GFP | Kind gift from Prof. Lenka Libusova, Charles University |
| GFP-SWIP | Kind gift from Prof. Lenka Libusova, Charles University |
| GFP-WASH1 | Kind gift from Prof. Alexis Gautreau, CNRS |
| VPS35-GFP | Kind gift from Prof. Peter J. Cullen, University of Bristol |
| VPS26-GFP | PCR amplified from cDNA and inserted into EcoR1/BamH1 in pEGFP-N1 |
| GFP-VPS29 | Kind gift from Prof. Peter J. Cullen, University of Bristol |
| TRIM62-pGAD | TRIM62 was PCR amplified from pTRIM62-GFP and inserted into pGADT7-AD (pGADT7-AD was a kind gift from Dr. Mahak Sharma) |
| WASH1-pGBK | WASH1 was PCR amplified from pGFP-WASH1 and inserted into pGBKT7 (pGBKT7 was a kind gift from Dr. Mahak Sharma) |
| CCDC53-pGBK | CCDC53 was PCR amplified from pCCDC53-GFP and inserted into pGBKT7 |

| | |
|---|---|
| FAM21-pGBK | FAM21 was PCR amplified from pFAM21-GFP and inserted into pGBKT7 |
| SWIP-pGBK | SWIP was PCR amplified from pSWIP-GFP and inserted into pGBKT7 |
| Strumpellin-pGBK | Strumpellin was PCR amplified from pStrumpellin-GFP and inserted into pGBKT7 |
| 3xFLAG | Kind gift from Dr. Santosh Chauhan, CCMB |
| FAM21-FLAG | FAM21 was PCR amplified from pFAM21-GFP and inserted into Xba1/Sal1 in pLenti CMV GFP Puro (Addgene #17448). FLAG sequence was added in primer |
| FAM21-FLAG mutants | Sequences amplified from pFAM21-GFP and inserted into Xba1/Sal1 in pLenti CMV GFP Puro. FLAG sequence was added in primer |
| gTRIM62-PX459 | gRNA was assembled and cloned in pSpCas9(BB)-2A-Puro (PX459) (PX459 is a kind gift from Dr. Debojyoti Chakraborty, IGIB) |
| CMVR8.74 | Banerjee lab |
| MD2.G | Banerjee lab |
| RFP-Rab7 | Banerjee lab |

**Supplementary Table 2. List of siRNAs used in this study**

| Gene | siRNA sequence |
|---|---|
| siTRIM62 #1 | CCGCGAGAAGTTCCCTGGCAA |
| siTRIM62 #2 | **CTGGTTTAATTAGACAAGGAT** |
| siTRIM62 #3 | CTGACCGACATCGAGCAGAAA |
| siWASH1 #1 | TGGGAACTAAGCAGATCACAA |
| siWASH1 #2 | **CAGCTCCTTGCTGCTCTTCAA** |
| siCCDC53 #1 | ATGGTTCAAGTGGGTGTACCA |
| siCCDC53 #2 | **AAGATATGCCAGATATCTCAA** |
| siSWIP #1 | **TCGGGCTTACGTAACTCACTA** |
| siSWIP #2 | AAGGTACCAGCCATCACTCTA |
| siStrumpellin #1 | **CAGAGAGTGCCTATCAACGAA** |
| siStrumpellin #2 | CAACGTGGAGCAAGAGTGTAA |
| siFAM21 #1 | AAGGTGAGAGATTATACTTCA |
| siFAM21 #2 | **ATGAGGAGGACCAGTGGAATA** |
| siVPS35 #1 | CACCATACTCCTTTCCATGTA |
| siVPS35 #2 | **CAGAATTGCCCTTAAGACTTT** |
| siACTR3 | CCGGCTGAAATTAAGTGAGGA |
| siWAVE | TGGTTTGGTAATGATCTGCAA |
| siN-WASP | CAGATACGACAGGGTATCCAA |
| siTRIM27 | GAACCAGCTCGACCATTTATT |
| siCAV1 | AACTAAACACCTCAACGATGA |
| siCLTC | TAATCCAATTCGAAGACCAAT |
| siATP6V1B2 | **CACGGTTAATGAAGTCTGCTA** |

*Bold indicates the siRNA used in the IAV entry experiments

**Supplementary Table 3. List of RT-PCR primers used in this study**

| Gene | | Primer Sequence |
|---|---|---|
| *TRIM62* | Forward | TACTGGGAGGTGGTGGTGGC |
| | Reverse | GTCATCAGCATTGTAGAAGATG |
| *WASH1* | Forward | AGAAGCAGCAGCAGAAGGAG |
| | Reverse | TACGATTCCCAGTCGTCCTC |
| *CCDC53* | Forward | GAAGCCACTTCAGAGCAACC |
| | Reverse | AGCATCTGGCCTCTCAAGAA |
| *SWIP* | Forward | TGCATGCAAGGCATTTAGAG |
| | Reverse | TTGAAAGGGTTTCGGTCATC |
| *Strumpellin* | Forward | CGAGGTTGCCAAGCTTACTC |
| | Reverse | GTCGCTCCCAACTCTTTCAG |
| *FAM21* | Forward | GGATCGGGAAGATACAAGCA |
| | Reverse | TCATCTTGACACGGCTCTTG |
| *VPS35* | Forward | CGTGAAGATGGACCTGGAAT |
| | Reverse | TCCACACGATCAGGGTAACA |
